# *Mycobacterium tuberculosis* DosS binds H_2_S through its Fe^3+^ heme iron to regulate the Dos dormancy regulon

**DOI:** 10.1101/2021.06.21.449194

**Authors:** Ritesh R. Sevalkar, Joel N. Glasgow, Martín Pettinati, Marcelo A. Marti, Vineel P. Reddy, Swati Basu, Elmira Alipour, Daniel B. Kim-Shapiro, Dario A. Estrin, Jack R. Lancaster, Adrie J.C. Steyn

**Affiliations:** Department of Microbiology, and Centers for AIDS Research and Free Radical Biology, University of Alabama at Birmingham, Birmingham, AL; Universidad de Buenos Aires, Facultad de Ciencias Exactas y Naturales, Departamento de Química Inorgánica, Analítica y Química Física, Buenos Aires, Argentina; CONICET-Universidad de Buenos Aires, Instituto de Química Física de los Materiales, Medio Ambiente y Energía (INQUIMAE), Buenos Aires, Argentina; Universidad de Buenos Aires, Facultad de Ciencias Exactas y Naturales, Departamento de Química Biológica, Buenos Aires, Argentina; CONICET-Universidad de Buenos Aires, Instituto de Química Biológica (IQUIBICEN), Buenos Aires, Argentina; Department of Physics, Wake Forest University, Winston-Salem, NC; Department of Pharmacology & Chemical Biology, Vascular Medicine Institute, University of Pittsburgh School of Medicine, Pittsburgh, PA; Africa Health Research Institute, University of KwaZulu-Natal, Durban, South Africa

**Keywords:** hydrogen sulfide, tuberculosis, dormancy regulon, two-component system, DosS

## Abstract

*Mycobacterium tuberculosis* (*Mtb*) senses and responds to host-derived gasotransmitters NO and CO via heme-containing sensor kinases DosS and DosT and the response regulator DosR. Hydrogen sulfide (H_2_S) is an important signaling molecule in mammals, but its role in *Mtb* physiology is unclear. We have previously shown that exogenous H_2_S can modulate expression of genes in the Dos dormancy regulon via an unknown mechanism(s). Here, we tested the hypothesis that *Mtb* senses and responds to H_2_S via the DosS/T/R system. Using UV-Vis and EPR spectroscopy, we show that H_2_S binds directly to the ferric (Fe^3+^) heme of DosS (K_D_ = 5.64 µM) but not the ferrous (Fe^2+^) form. No interaction with DosT was detected. Thus, the mechanism by which DosS senses H_2_S is different from that for sensing NO and CO, which bind only the ferrous forms of DosS and DosT. Steered Molecular Dynamics simulations show that H_2_S, and not the charged HS^-^ species, can enter the DosS heme pocket. We also show that H_2_S increases DosS autokinase activity and subsequent phosphorylation of DosR, and H_2_S-mediated increases in Dos regulon gene expression is lost in *Mtb* lacking DosS. Finally, we demonstrate that physiological levels of H_2_S in macrophages can induce Dos regulon genes via DosS. Overall, these data reveal a novel mechanism whereby *Mtb* senses and responds to a third host gasotransmitter, H_2_S, via DosS-Fe3^+^. These findings highlight the remarkable plasticity of DosS and establish a new paradigm for how bacteria can sense multiple gasotransmitters through a single heme sensor kinase.

**Significance Statement:** Hydrogen sulfide (H_2_S) is an important signaling molecule in eukaryotes and bacteria, and along with CO and NO, is an important part of host defense against *Mycobacterium tuberculosis* (*Mtb*). However, the mechanism(s) by which *Mtb* senses and responds to H_2_S is unknown. Here, we report that the *Mtb* heme sensor kinase DosS, a known sensor of CO and NO, is also a sensor of H_2_S. We found that H_2_S binds DosS in its ferric (Fe^3+^) state, which is considered as its inactive state, to induce the Dos dormancy regulon during infection. These data highlight the unusual capacity of *Mtb* to sense multiple gasotransmitters through a single sensing protein.

## Introduction

Tuberculosis (TB) is a global epidemic responsible for ∼1.4 million deaths annually (1). *Mycobacterium tuberculosis* (*Mtb*), the causal agent of TB, can persist in a state of clinical latency for decades. *Mtb* survives and establishes an infection due, in part, to its ability to sense and respond to host defenses in the lung, including host-generated gasotransmitters. Carbon monoxide (CO) and nitric oxide (NO) are critical components of the host defense to clear the pathogen and are important to the outcome of *Mtb* infection (2, 3). The most recent addition to the list of gasotransmitters is hydrogen sulfide (H_2_S). Notably, enzymes required for the generation of NO, (nitric oxide synthase [iNOS]) (3), CO (heme oxygenase-1 [HO-1]) (2, 4, 5), and H_2_S (cystathionine β-synthase [CBS] (6), cystathionine γ-lyase [CSE] (6), and 3-mercaptopyruvate sulfur transferase [3-MPST]) (6), are upregulated in the lungs of *Mtb*-infected mice and human TB patients. The increased levels of these enzymes suggest an abundance of NO, CO, and H_2_S at the primary site of infection.

The role of NO and CO in TB pathogenesis is well studied compared to that of H_2_S. We have recently shown that *Mtb*-infected mice deficient in the H_2_S-producing enzyme CBS (6, 7) or CSE (6, 7) survive significantly longer with reduced organ burden, suggesting that host-generated H_2_S is beneficial for *Mtb in vivo*. Bacterial two-component regulatory systems sense changes in the host environment and mediate adaptive genetic responses. The *Mtb* genome encodes 11 paired two-component systems (8), including the DosS/T-DosR system, comprised of heme-containing sensor kinases DosS and DosT and their cognate transcriptional response regulator DosR (9). DosS, currently regarded as a redox sensor, is considered inactive in the oxidized/met (Fe^3+^) state and is activated by direct binding of NO or CO to the heme iron in the reduced (Fe^2+^) state (10). DosT is an oxygen sensor and is inactive in its oxy-bound form which is activated upon loss of O_2_ or direct binding of NO or CO to the heme iron in the reduced (Fe^2+^) state (10, 11) (Fig. 1*A*).

**Fig. 1.**
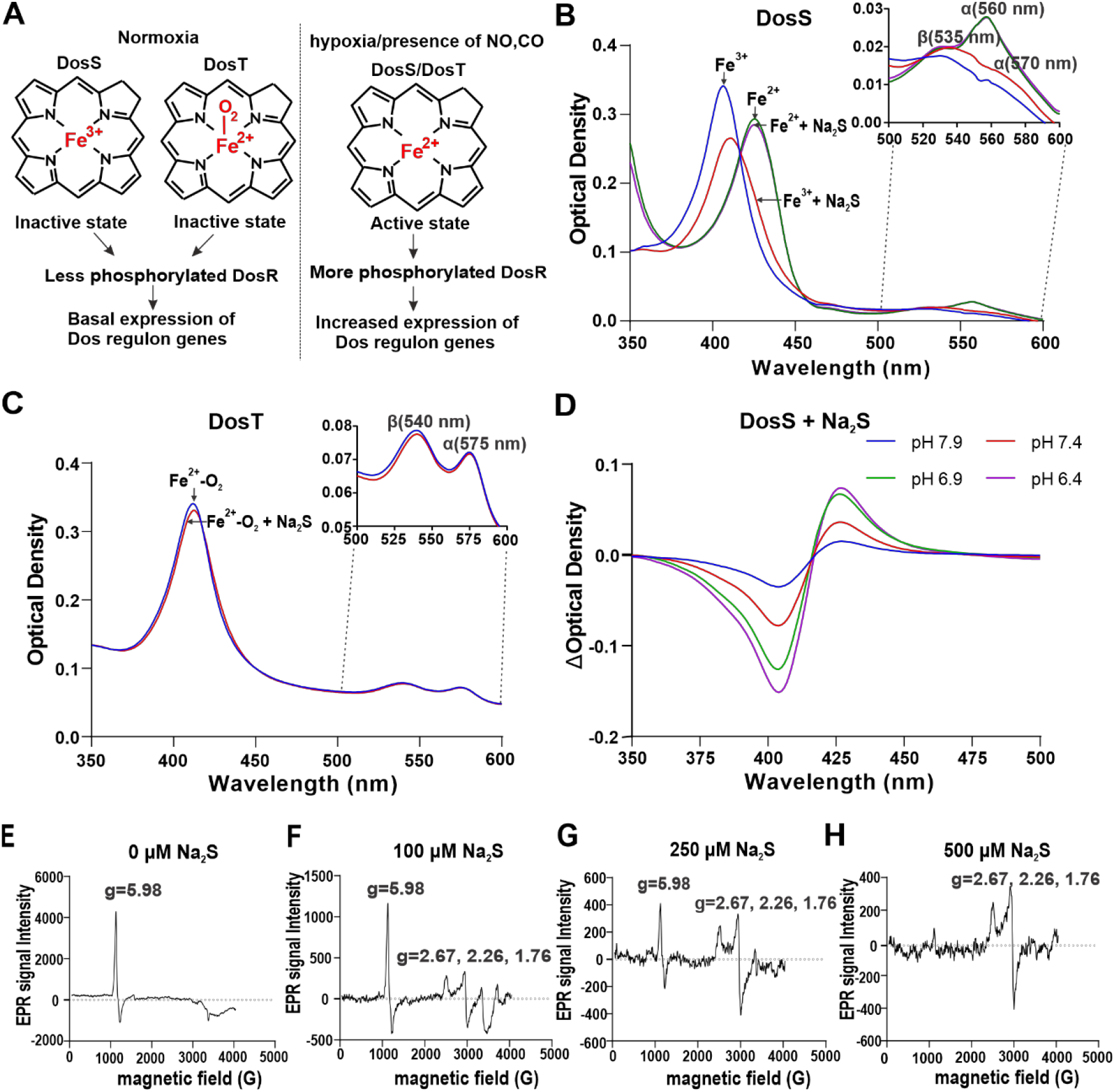
Characterization of DosS binding to H_2_S. (*A*) Depictions of DosS and DosT sensing of O_2_, NO and CO under various conditions. (*B*) Representative UV-Visible absorption spectra of recombinant DosS (3 µM) in the Fe^3+^ form (blue curve), the Fe^3+^ form in the presence of 100 µM Na_2_S (red curve), the Fe^2+^ form in the presence of 100 µM DTH (green curve) and the Fe^2+^ form in the presence of 100 µM DTH and 100 µM Na_2_S (orange curve). (*Inset*) Absorption spectra replotted to highlight the α and β absorption peaks of the Fe^3+^ form (α at 570 nm and β at 535 nm), and the Fe^2+^ form (α at 560 nm). (*C*) Representative UV-Visible absorption spectra of recombinant DosT (3 µM) in the Fe^2+^-O_2_ form (blue curve) and the Fe^2+^-O_2_ form in the presence of 100 µM Na_2_S (red curve). (*Inset*) Absorption spectra replotted to highlight the α (570 nm) and β (535 nm) peaks. (*D*) Changes to the UV-Visible absorption spectra of the Fe^3+^ form of recombinant DosS (3 µM) resulting from the addition of 25 µM Na_2_S at different pH conditions, relative to DosS without Na_2_S. (*E-H*) EPR spectroscopic analysis of recombinant DosS showing a single axial peak characteristic of paramagnetic high-spin Fe^3+^ heme iron. DosS alone (*E*), and in the presence of increasing concentrations of Na_2_S (*F-H*) which shows the appearance of additional peaks characteristic of the low-spin state of heme iron. UV-Vis spectra are representative of at least 5 independent measurements.

We recently reported that exposure of *Mtb* to the H_2_S donor GYY4137 induces the expression of genes that regulate cysteine metabolism and genes in the copper and Dos dormancy regulons (7). While H_2_S can chemically modify biomolecules directly (12-14), we considered the possibility that alterations in gene expression in response to H_2_S are mediated by regulatory proteins in *Mtb*. Since H_2_S is known to bind the iron in heme-containing proteins (15-19), and because DosS was among the Dos regulon genes induced upon exposure to H_2_S (7), we hypothesize that DosS and/or DosT sense and respond to H_2_S to induce the Dos dormancy regulon. To examine the interaction of H_2_S with DosS and DosT, we used UV-visible and EPR spectroscopy. We also examined how H_2_S modulates DosS autokinase and phosphate transfer, and *Mtb* gene expression *in vitro*. Lastly, we examined how *Mtb* senses H_2_S during macrophage infection. We expect that the findings in this work will lead to an improved understanding of *Mtb* persistence.

## Results

### DosS in the Fe^3+^ form binds H_2_S

DosS and DosT contain heme and exhibit specific absorption characteristics in the ultraviolet-visible (UV-Vis) range, which are altered upon interaction between the heme iron and ligands like NO and CO (10, 20). To determine whether Mtb DosS or DosT sense H_2_S via its heme iron, we monitored spectral changes of the ferric (Fe^3+^) and ferrous (Fe^2+^) forms of recombinant DosS and DosT in the presence of sulfide (here, we define sulfide as H_2_S and HS^-^) following addition of sodium sulfide (Na_2_S). Addition of sulfide red shifts the Soret peak of the Fe^3+^ form of DosS from 408 nm to 415 nm, indicative of a high-spin to low-spin transition (21), with increased peak intensities of the α (570 nm) and β (535 nm) bands (Fig. 1*B*), similar to spectral changes observed upon sulfide binding in other heme-containing proteins (Table1). Reduction of DosS to the Fe^2+^ form using sodium dithionite (DTH) shifts the Soret peak to ∼425 nm with emergence of a new peak at 560 nm, as observed previously (10, 22). However, the absorption pattern of DTH-reduced DosS remains unchanged in the presence of sulfide. Similarly, addition of sulfide does not alter the absorption spectrum of DosT, where the heme iron remains in the oxy-bound state (Fe^2+^-O_2_) (Fig. 1*C*) (10, 22). These data suggest that sulfide directly interacts with the Fe^3+^ form of DosS. This is mechanistically distinct from the binding of NO and CO, which bind the heme iron of DosS and DosT in the Fe^2+^ state.

H_2_S is in protonation equilibrium with HS^-^ in solution with a pKa value of 7.01 (23). To determine whether H_2_S or anionic hydrosulfide (HS^-^) binds to the Fe^3+^ form of DosS, we monitored the UV-Vis spectra of DosS at pH values above and below the pKa of H_2_S in the presence of 25 µM Na_2_S. Notably, as the pH decreased we observed lower peak intensities in the 405 nm range, indicating reduced levels of unbound DosS, and increases in the ∼420 nm range corresponding to sulfide-bound DosS. These data suggest that H_2_S, and not HS^-^, is the sulfide species that initially binds to the heme iron of DosS (Fig. 1*D*).

Iron is paramagnetic in its Fe^3+^ state. Ligand binding to the heme iron results in perturbations in the *d*-orbitals that can be monitored by electron paramagnetic resonance (EPR) spectroscopy (10, 24). To confirm that H_2_S binds directly to the Fe^3+^ form of DosS, we compared the EPR spectra of DosS before and after exposure to sulfide. DosS alone gives a strong axial feature centered at g=5.98, which is indicative of Fe^3+^ in the high-spin (S=5/2) state (Fig. 1*E*) (10, 25) (Table 1). The addition of increasing concentrations of sulfide results in the conversion of the g=5.98 high-spin signal into a rhombic low-spin signal with g values of 2.67, 2.26, and 1.76 (Fig. 1 *F*-*H*), which are similar to low-spin sulfide-bound species reported for other heme-containing proteins (Table 1). Taken together, these results indicate that H_2_S is a ligand of the Fe^3+^ form of DosS and binding of H_2_S converts the Fe^3+^ heme iron from the high-spin state to low-spin state.

**Table 1:** Comparison of Biophysical Parameters of Sulfide-Binding Proteins.

| Protein | UV-vis (nm) | EPR (g-values) | K<sub>D</sub> (μM) | Reference |
| --- | --- | --- | --- | --- |
| *Mtb* DosS | 415, 535, 570 (s) | 2.67, 2.26, 1.76 | 5.64 (H<sub>2</sub>S) | This work |
| *L. pectinata* Hb I | 425, 545, 573 (s) | 2.67, 2.24, 1.84 | 0.09 (XS<sub>T</sub>) | (1) |
| Human Hemoglobin | 423, 577, 541 | 2.51, 2.25, 1.86 | 17 (XS<sub>T</sub>) | (2) |
| Bovine Hemoglobin | 425, 542, 575(s) | 2.55, 2.26, 1.88 | 7(XS<sub>T</sub>) | (3) |
| Equine Myoglobin | 427, 547, 580 | 2.56, 2.25, 1.83 | 96 (XS<sub>T</sub>) | (4), (5) |
| Human Myeloperoxidase | 432, 625 | 2.567, 2.274, 1.850;<br>2.512, 2.262, 1.875 | < 12 (XS<sub>T</sub>) | (6) |
| Microperoxidase | 414, 536, 568 | -- | 219 (calc) (XS<sub>T</sub>) | (7) |
| Truncated *B. subtilis* Hb | 427, 550, 577 | -- | 0.2 (XS<sub>T</sub>) | (8) |
| Truncated *T. fusca* Hb | 425, 550, 575 | -- | 0.36 (XS<sub>T</sub>) | (8) |
| *E. coli* DOS (heme-regulated TCS PDE) | 427, 546, 579 | -- | ca. 200 (XS<sub>T</sub>) (based on PDE activation) | (9) |
| *Anaeromyxobacter* AfGCHK (heme-regulated TCS His kinase) | 423-426, 549-551 | -- | 200-5000 (XS<sub>T</sub>) for binding, although no autokinase activation with 10 mM | (10) |

Note: Hemeproteins that get reduced on sulfide binding do not show the EPR spectrum and hence were not used for comparison eg., cytochrome c oxidase, neuroglobin, cytochrome c.
H<sub>2</sub>S = based on H<sub>2</sub>S concentration
XS<sub>T</sub> = based on concentration of total sulfide species (H<sub>2</sub>S + HS<sup>-</sup> + S<sup>2-</sup>)
TCS = two-component system
(s) = shoulder
calc = calculated

### Molecular dynamics simulations show that H_2_S, but not HS^-^, enters the DosS heme pocket

The DosS heme group is buried within a hydrophobic pocket that is enclosed within the N-terminal GAF-A domain (22, 26). Access to the hydrophobic heme pocket is limited, and ligand entry is influenced by the adjacent amino acid side chains (22). Steered Molecular Dynamics (sMD) has been used to estimate association free energy required for H_2_S and HS^-^ to access the heme iron in *L. Pectinata* met-hemoglobin (27) and met-myoglobin (27, 28). Similarly, our sMD simulations estimate the free energy barriers for access to the DosS heme iron to be approximately 5.6 kcal/mol for H_2_S and 16.7 kcal/mol for HS^-^ (Fig. 2*A*). The much higher free energy barrier for HS^-^ indicates that the uncharged H_2_S species is strongly favored to enter the heme pocket.

**Fig. 2.**
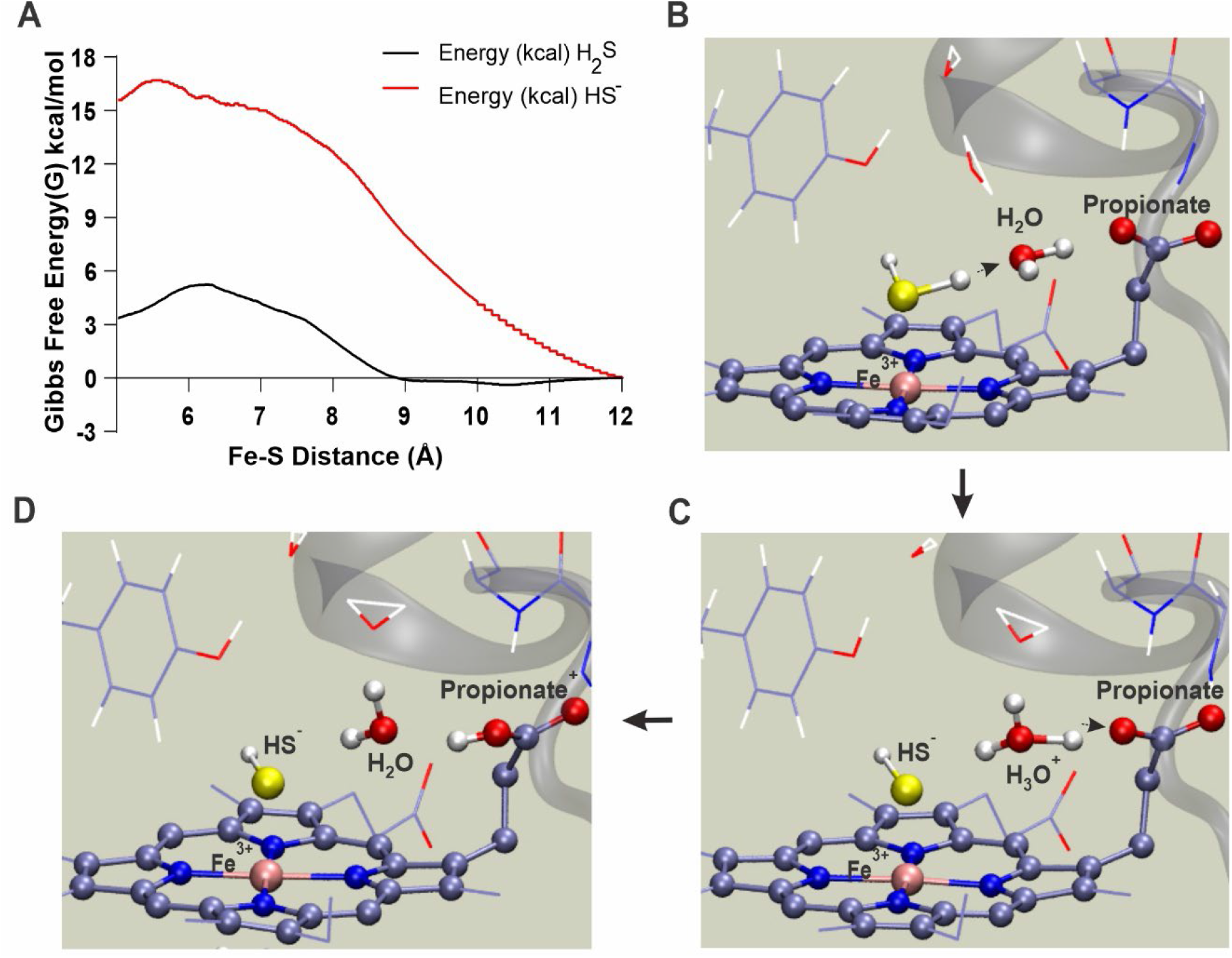
Molecular modeling of DosS interaction with H_2_S. (*A*) Calculated average association free energy profiles for H_2_S and HS^-^ as a function of intermolecular distance between the DosS heme iron and sulfur. Free energy values were generated by employing 98 separate trajectories for H_2_S and 89 trajectories for HS^-^ using a steered Molecular Dynamics (sMD) approach. (*B-D*) QM/MM MD simulation snapshots depicting the steps of a predicted proton transfer from H_2_S to a nearby heme propionate group through a water bridge within 0.3 ps of MD. The QM subsystem atoms are shown in ball and stick representation.

Our sMD simulations predict that H_2_S accesses the heme iron by passing between Phe^98^ and Leu^114^ of the heme binding pocket (Fig. S1). These residues form a “gate” which when open allows H_2_S and a few water molecules into and out of the heme pocket. This is possible due to the neutral charge of H_2_S (Fig. S1 *A-C*, Movie S1). In contrast, our sMD simulations indicate that HS^-^ carries a strongly-bound solvation sphere that pushes the Phe^98^ and Leu^114^ side chains away, resulting in a considerable number of water molecules entering the heme pocket and increasing solvation of the heme active site (Fig. S1 *D-F*, Movie S2). The process of HS^-^ entry is energetically unfavorable, as indicated by a much higher predicted free energy barrier compared to H_2_S. Notably, this H_2_S entry “gate” is located away from the “water channel” identified by Cho, et. al., in the DosS GAF-A domain structure (22). Overall, our sMD modeling indicating more favorable heme access for H_2_S supports our UV-Vis data demonstrating increased binding of sulfide at lower pH (Fig. 1*D*).

### Quantum mechanics simulations show that H_2_S deprotonates following heme iron binding

Modeling studies of heme-containing proteins predict that H_2_S can deprotonate following binding to heme iron (27, 28). To characterize the heme-bound state of H_2_S in DosS, we employed combined quantum mechanics/molecular mechanics simulations with density functional theory calculations (QM[DFT]/MM). Table 2 shows the structural and electronic parameters of both H_2_S and HS^-^ bound states of DosS. As expected, the ligand-bound states display a low-spin ground state. The structural analysis shows that HS^-^ forms a tighter bond (a significantly smaller Fe-S distance, [d Fe-S = 2.17 Å]) due to a significant charge transfer (sigma donation). Further, the proximal Fe-His bond shows a slightly positive trans effect, displaying a slightly smaller distance compared to penta-coordinated heme (ca 2.12 Å) (29, 30). Interestingly, in the H_2_S-bound state, the two protons become asymmetric and a weakening of the S-H bond is observed (d Fe-S = 2.35 Å). On this basis, we evaluated the possibility of deprotonation of Fe-bound H_2_S using hybrid QM/MM simulations. Strikingly, a simulation duration of 1 picosecond (ps) was sufficient to observe deprotonation of Fe-bound H_2_S (Fig. 2*B-D* and Movie S3). The proton acceptor is a water molecule located opposite from the distal Tyr (i.e., close to the solvent-exposed heme edge) which subsequently transfers a proton to the heme propionate (Fig. 2*D*). The heme propionate is accessible to the solvent and can again be deprotonated or remain in a protonated state. Overall, these data strongly suggest that HS^-^ is the tighter binding and predominant bound sulfur species.

**Table 2.**
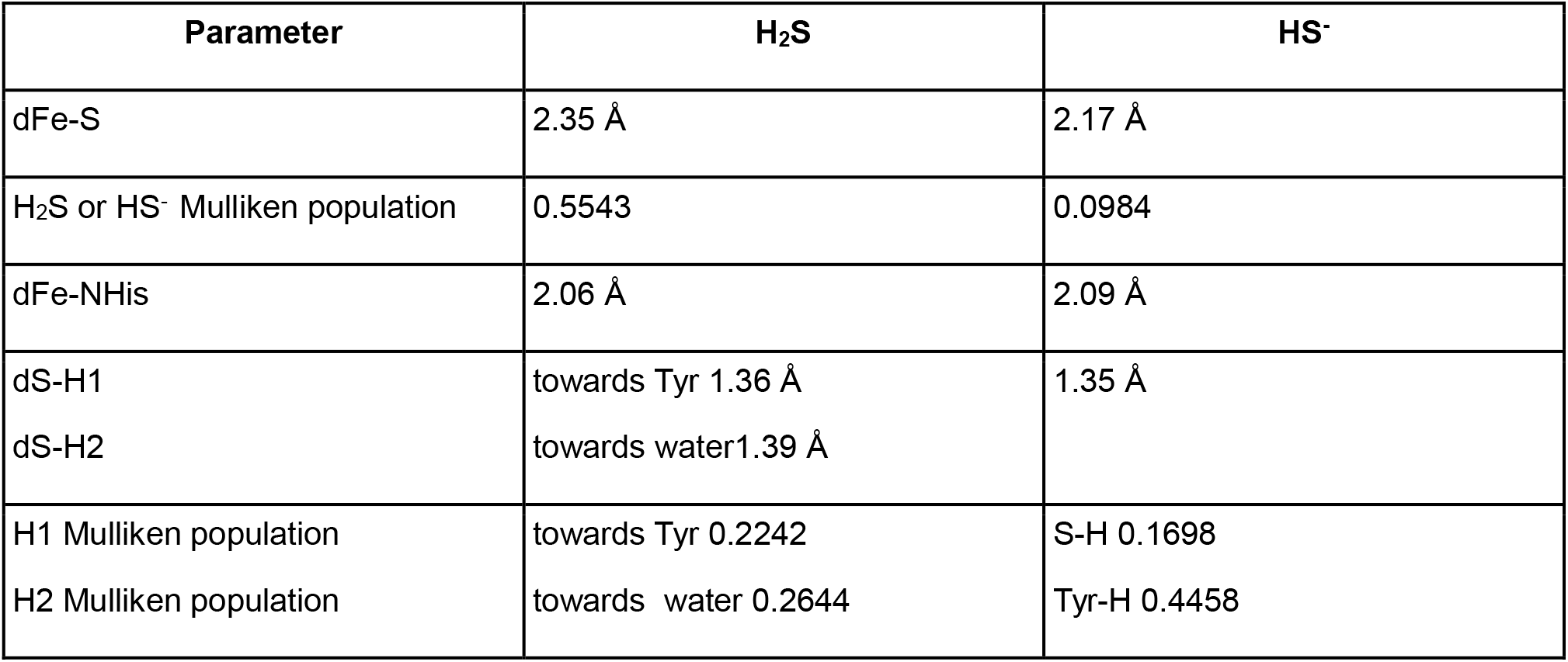
Comparison of structural and electronic parameters of H2S and HS^-^ bound states obtained from QM/MM calculations.

### H_2_S does not reduce the heme iron in DosS and binds with low micromolar affinity

H_2_S is known to reduce heme iron to the Fe^2+^ state in proteins like myoglobin and hemoglobin (16). Since the Fe^2+^ form of DosS is understood to be the active form of the kinase (10, 20), it is important to determine whether H_2_S binding reduces the DosS heme iron. Therefore, we monitored the UV-Vis spectrum of the Fe^3+^ form of DosS in the presence of 200 µM Na_2_S over time. As shown in Fig. 3*A*, addition of Na_2_S resulted in the expected shift in the Soret peak from 408 nm to 415 with increased peak intensity of the α (570 nm) and β (535 nm) bands. Notably, we observed no additional shift in the Soret peak to 425 nm nor the appearance of the peak at 560 nm, both of which are indicative of the reduced form of DosS (Fig. 1*A*). Over the course of the experiment we noted a decrease in overall absorbance, which is most likely due to the loss of the heme prosthetic group from DosS (31, 32). These results indicate that H_2_S binding does not readily alter the oxidation state of the DosS heme iron.

**Fig. 3.**
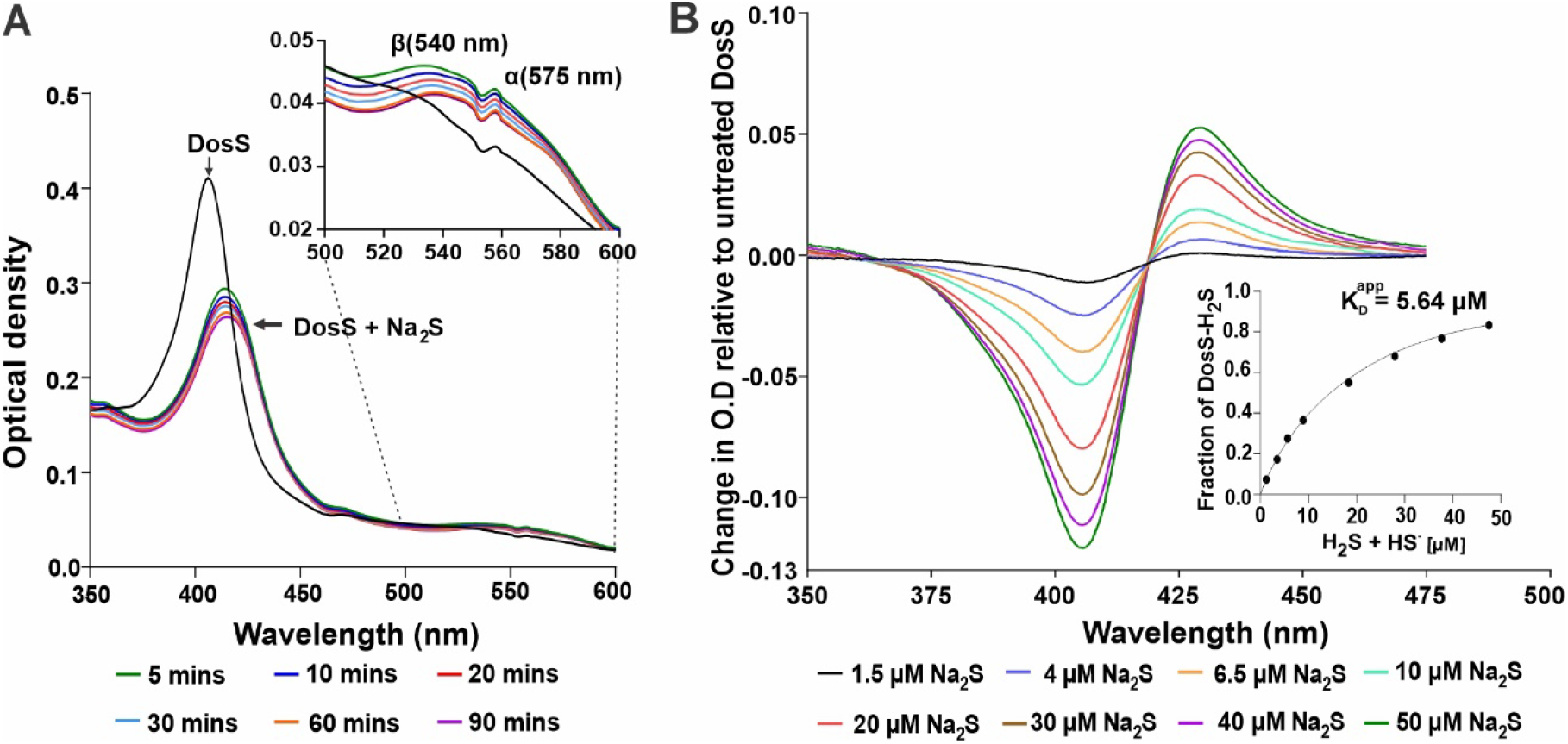
H_2_S does not reduce the DosS heme iron and binds with low micromolar affinity. (*A*) Representative UV-Visible spectra of the Fe^3+^ form of DosS (3 µM) for 5-90 minutes following the addition of 200 µM Na_2_S. (*Inset*) Absorption spectra replotted to highlight the α (570 nm) and β (535 nm) peaks. The lack of red shift over time following addition of Na_2_S as well as the absence of a strong α peak at 560 nm indicate that the DosS heme iron is not reduced in the presence of sulfide within 90 minutes. A small spectral artifact caused by spectrophotometer filter switching is present at ∼550 nm. (*B*) Representative changes to the UV-Visible spectra of recombinant DosS (3 µM) resulting from the addition of different concentrations of Na_2_S, relative to DosS without Na_2_S. (*Inset*) Substrate saturation curve of DosS-H_2_S binding was generated using UV-Vis absorption data points obtained from the titration DosS with increasing concentrations of Na_2_S. UV-Vis spectra are representative of at least 3 independent assays.

To determine the affinity of DosS for H_2_S, we next monitored changes in the absorbance spectrum of DosS in the Fe^3+^ state over a wide range of Na_2_S concentrations (Fig. 3*B*). Using the maximum change in absorbance at ∼408 nm, we generated a substrate saturation curve from which a K_D_^app^ value of 5.64 ± 0.27 µM was calculated for H_2_S binding to DosS (3*B* and *Inset*). Notably, this K_D_ ^app^>value is similar to the K_D_^app^ value of 7.0 ± 0.4 µM reported for the H_2_S-met hemoglobin complex (15, 18). In summary, these results show that measureable, direct binding of H_2_S to DosS (Fe^3+^) does not reduce DosS to the active Fe^2+^ form.

### H_2_S increases DosS autokinase activity and DosR phosphorylation

To address the question of whether H_2_S binding can alter DosS kinase activity, we performed kinase assays using γ-^32^P-labeled ATP and recombinant DosS. We determined relative autokinase activities of DosS (Fe^2+^), DosS (Fe^3+^), and DosS (Fe^3+^-HS^-^) at 5-60 minutes following addition of γ-^32^P labeled ATP. Compared to the Fe^3+^ form of DosS, which is considered the least active form (10), the Fe^3+^-HS^-^ form of DosS exhibited increased autokinase activity. In these assays, the Fe^2+^ form of DosS showed the highest autokinase activity (Fig 4*A*). Densitometric analysis of autokinase radiograms shows that the activity of the Fe^3+^-HS^-^ form of DosS is increased by an average of ∼30% at all time points compared to unbound DosS in the (Fe^3+^) form (Fig. 4*B*).

**Fig. 4.**
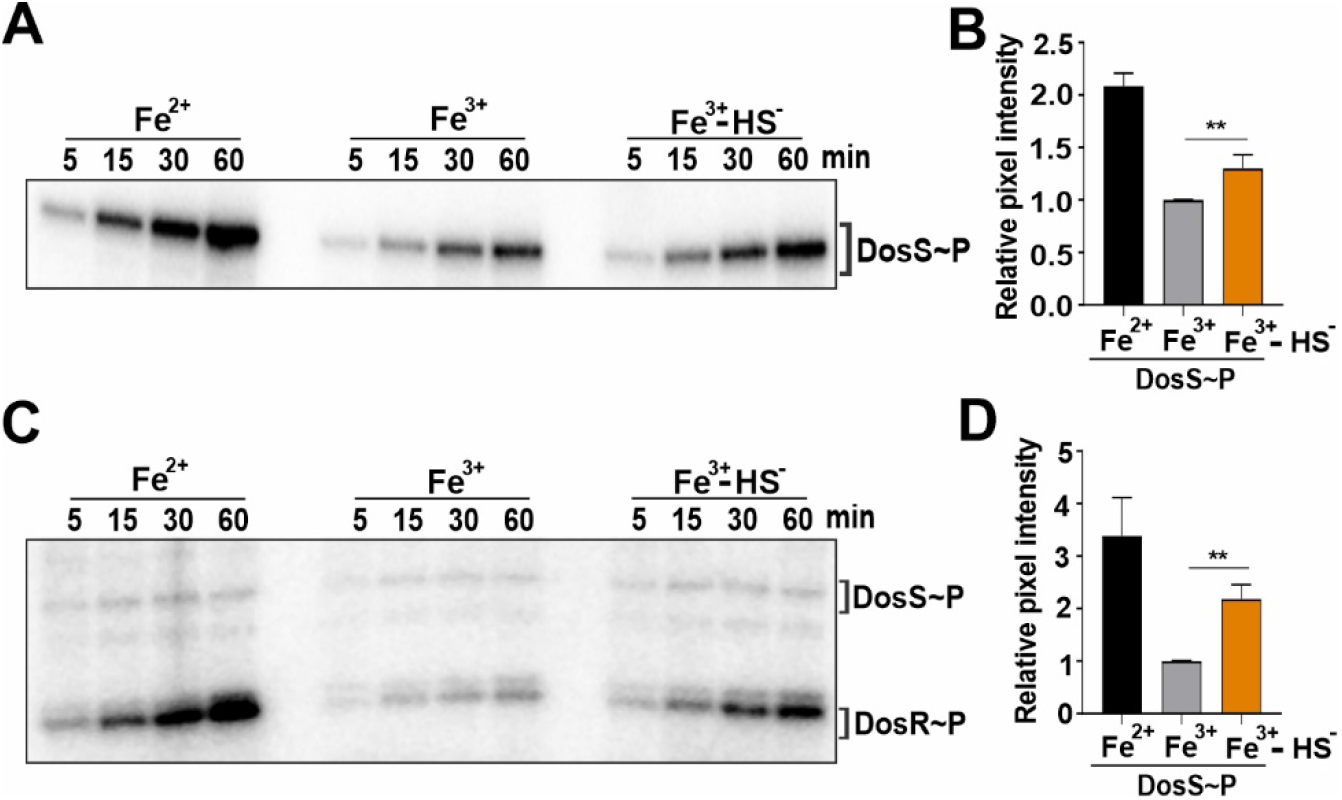
H_2_S stimulates DosS and DosR phosphorylation. (*A*) Representative autoradiogram of PAGE-resolved ^32^P-labeled recombinant DosS following autophosphorylation in the presence of γ-^32^P-ATP alone (left and center panels) or with γ-^32^P-ATP in the presence of 100 µM Na_2_S (right panel). (*B*) Densitometric quantitation of DosS bands in (*A*) at 60 min (n=3). (*C*) Representative autoradiogram of PAGE-resolved recombinant DosS and DosR following phosphorylation of DosR by γ-^32^P-labeled DosS alone (left and center panels) or with γ-^32^P-labeled DosS in the presence of 100 µM Na_2_S (right panel). These reactions were performed by adding γ-^32^P-labeled ATP to a reaction containing both DosS and DosR and analyzed at different time points. (*D*) Densitometric quantitation of DosR bands in (*C*) at 60 min (n=3). Data in (*B*) and (*D*) are shown as the mean ±SEM and were analyzed using one-way ANOVA with Tukey’s multiple comparisons test performed using GraphPad Prism version 9. **P < 0.01. Autoradiograms are representative of at least 3 independent assays.

Next, we sought to determine whether the H_2_S-mediated increase in DosS autokinase activity results in augmented phosphorylation of DosR, the cognate response regular of DosS. First, to standardize our transphosphorylation assays, unphosphorylated recombinant DosR was added to reaction buffer containing γ-^32^P-labeled DosS. Surprisingly, the phosphorylation of DosR by labeled DosS was extremely rapid, and was completed within seconds (Fig. S2). Therefore, to observe differences in the rates of phosphate transfer of DosS (Fe^2+^), DosS (Fe^3+^), and DosS (Fe^3+^-HS^-^), unphosphorylated DosS and DosR were added to the reaction buffer prior to the addition of γ-^32^P-labeled ATP. Under these conditions, we observed increased phosphorylation of DosR in the presence of DosS (Fe^3+^-HS^-^) compared to DosR phosphorylation in the presence of DosS (Fe^3+^) at all time points measured. Maximum phosphorylation of DosR was observed in the presence of DosS (Fe^2+^) (Fig. 4*C*). Densitometric analysis of transphosphorylation radiograms shows an ∼2-fold increase in labeled DosR in the presence of DosS (Fe^3+^-HS^-^) compared to the Fe^3+^ form of DosS (Fig. 4*D*). Overall, these data indicate that the binding of sulfide to DosS increases its autokinase activity. Given the extremely rapid transfer of phosphate from DosS to DosR, we conclude that increased DosR phosphorylation is attributable primarily to changes in autokinase activity.

Our findings that sulfide binding increases autokinase activity and that HS^-^ is likely the predominant bound ligand have implications for the structural basis of kinase domain activation. To elucidate a mechanism by which sulfide binding increases DosS autokinase activity, we employed MD modeling to detect structural differences between DosS in the Fe^3+^ and Fe^3+^-HS^-^ forms. Our analysis indicates that the hydrogen bonding network distal to the heme is significantly different between the ferric high spin (off-state) and the Fe^3+^-HS^-^ low spin state (Fig. 5), but not the Fe^3+^-H_2_S state. In the Fe^3+^ state of DosS, there is a loosely coordinated water molecule which cannot act as a hydrogen bond acceptor for Tyr^171^. Therefore the hydroxyl group of Tyr^171^ rotates upward and instead forms a tight hydrogen bond with Glu^87^ (Fig. 5*A*). In contrast, when HS^-^ is bound to DosS in the Fe^3+^ state, Tyr^171^ hydroxyl group rotates downward and establishes a tight hydrogen bond with HS^-^ with the negative sulfur atom as the hydrogen bond acceptor. This results in the release of Glu^87^, which then moves closer to, and establishes a tight hydrogen bond with, His^89^ and Thr^169^ resulting in changes in their relative position, particularly for the loop in the peptide backbone in which H^89^ is located. Disruption of the hydrogen bond between Tyr^171^ and Glu^87^ is further promoted by the presence of water molecules between them (Fig 5*B*). These observations suggest that the DosS off-state is characterized by strong Tyr^171^-Glu^87^ interaction, while the on-state is characterized by a strong Tyr^171^-HS^-^ interaction that releases Glu^87^, which is consistent with the observation that CO/NO-bound DosS is active and shows the disruption of the hydrogen bonding network between Tyr^171^-Glu^87^-His^89^ (26). Overall it appears that signal transmission to the histidine kinase domain is initiated by disruption of Glu^87^-Tyr^171^ H-bonding and further amplified by positional changes in the His^89^-containing loop.

**Fig. 5.**
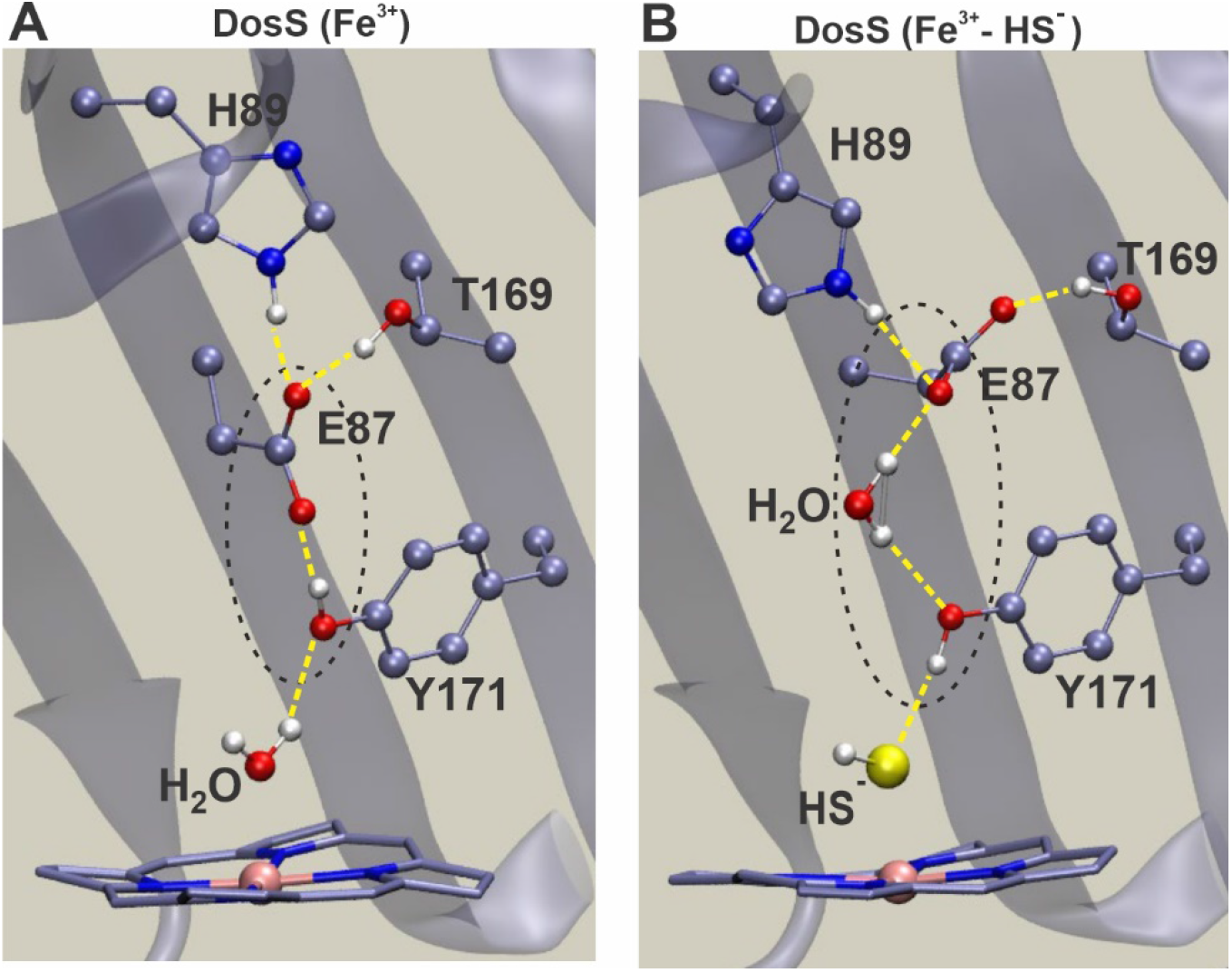
Sulfide binding alters the hydrogen bonding network in the DosS heme pocket. (*A*) Hydrogen bonding network in the distal domain of ferric (Fe^3+^) DosS showing an intact H-bond between glutamate (E87) and tyrosine (Y171). (*B*) MD-modeled hydrogen bonding network in the distal domain of ferric (Fe^3+^) DosS in the presence of sulfide showing disrupted H-bonding between glutamate (E87) and tyrosine (Y171). Predicted changes in the H-bonding patterns may lead to structural changes which alter DosS kinase activity upon sulfide binding.

### DosS senses H_2_S to regulate the Dos dormancy regulon

To determine whether the H_2_S-mediated activation of DosS is sufficient to drive increased expression of Dos regulon genes, we exposed WT *Mtb* and *Mtb* Δ*dosS*, a *dosS* deletion mutant (33), to Na_2_S. This was followed by quantitation of mRNA transcripts of representative Dos regulon genes. As shown in Fig. 6*A, fdxA, hspX, rv2030c* and *rv2626* transcript levels were markedly increased in WT *Mtb*, but not Δ*dosS*, following exposure to Na_2_S. These results indicate that H_2_S mediates increased expression of Dos regulon genes via DosS activation, consistent with our previous observations (7).

**Fig. 6.**
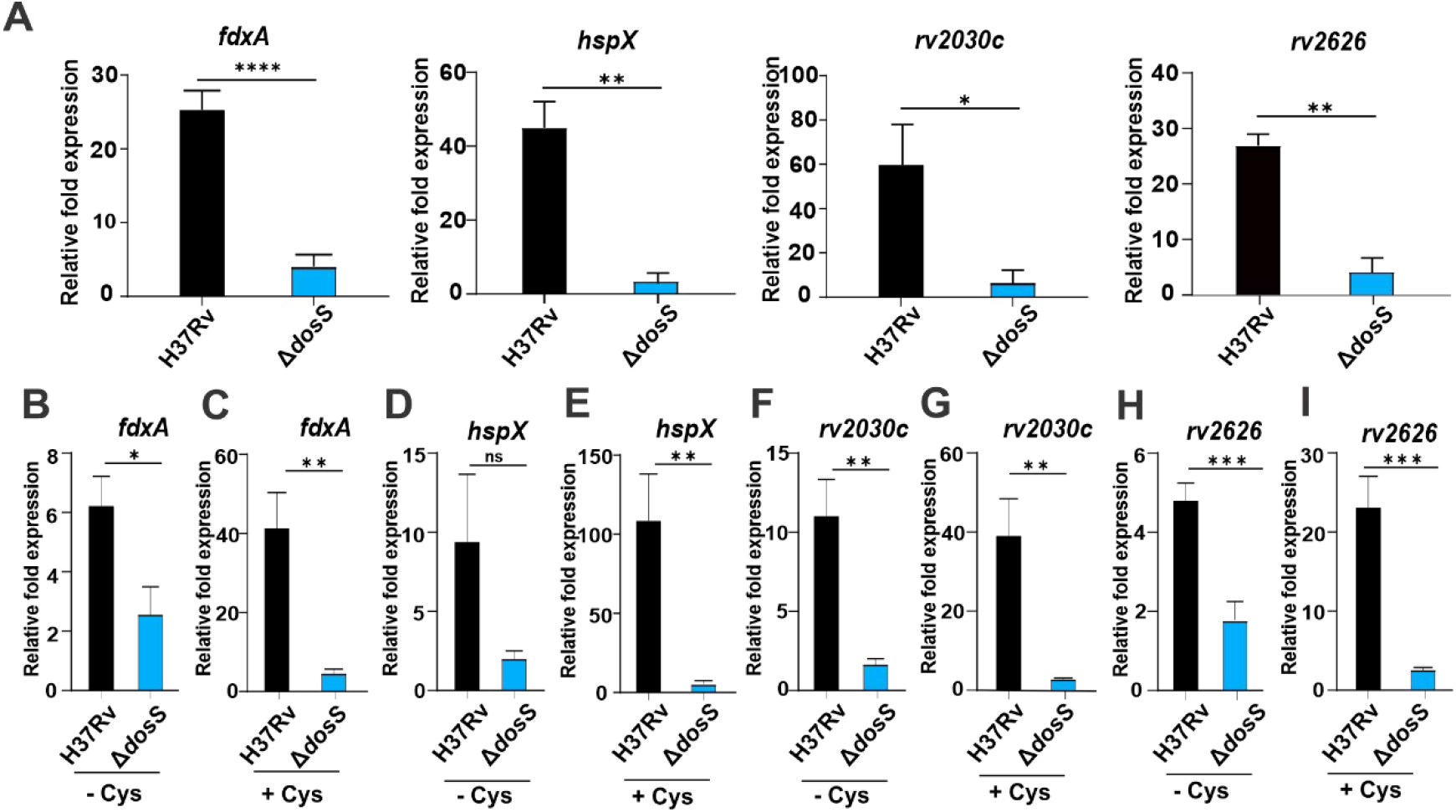
Effect of H_2_S on expression of DosR regulon genes. (A) Expression of representative Dos regulon genes in WT and ΔdosS *Mtb* cells exposed to 50 μM Na_2_S for 30 mins, relative to unexposed *Mtb* cells (n=6, 3 independent experiments with qRT-PCR performed in duplicate). (B-I) Expression of representative Dos regulon genes in WT and ΔdosS *Mtb* isolated from infected RAW 264.7 macrophages grown with or without 0.2 mM L-cysteine for 24 hours (n=6, 3 independent experiments with qRT-PCR performed in duplicate). RT-qPCR gene expression data from macrophage-isolated *Mtb* are shown relative to gene expression in WT and ΔdosS *Mtb* cultures exposed to DMEM. Data are shown as the mean ±SEM and were analyzed using a unpaired t-test performed using GraphPad Prism version 9. *q < 0.05, **q < 0.01, ***q < 0.001 and ****q < 0.0001.

We next sought to determine whether host-generated H_2_S can be sensed by DosS to mediate induction of Dos regulon genes. CBS and CSE can produce H_2_S using L-cysteine as a substrate (34, 35). To avoid potentially confounding effects of CBS/CSE enzyme inhibitors on *Mtb* as seen previously (7), we chose to use Cys to modulate intracellular levels of H_2_S in RAW 264.7 macrophages. First, we examined H_2_S levels in RAW 264.7 macrophages grown in media supplemented with L-cysteine using the fluorescent WSP-5 H_2_S-sensing probe (36). We observed an increased intracellular fluorescence signal in RAW 264.7 macrophages grown in media containing 0.1-2.0 mM L-cysteine compared to cysteine-free media, with maximum signal observed at 0.2 mM L-cysteine (Fig. S3). This finding demonstrates that addition of exogenous L-cysteine increases H_2_S production. Importantly, levels of Cys in serum and cells have been reported to range between 30-260 μM (37-40). On this basis, RAW 264.7 macrophages were grown in cysteine-free media or media containing 0.2 mM L-cysteine and infected with WT or Δ*dosS Mtb*. At 24 hours post-infection, intracellular bacteria were recovered and RNA was extracted. We observed marked increases in transcript levels of *fdxA, hspX, rv2030c*, and *rv2626* in WT *Mtb*, but not Δ*dosS Mtb*, isolated from macrophages grown in media containing 0.2 mM L-cysteine (Fig. 6*B-I*). These data indicate that *Mtb* senses host-derived H_2_S via DosS to induce the expression of Dos regulon genes.

## Discussion

The *Mtb* DosS/T/R signal transduction system is known to sense three host-derived dormancy signals, NO (10, 11, 33), O_2_ (33, 41), and CO (4, 5) to induce the 48-gene Dos dormancy regulon. Here, we report that *Mtb* is capable of sensing a fourth gasotransmitter, H_2_S, to induce the Dos dormancy regulon. Importantly, we show that DosS is capable of sensing H_2_S at levels produced by macrophages, resulting in the upregulation of key dormancy regulon genes. We also show that *Mtb* DosS, but not DosT, can sense H_2_S and does so via its heme iron in the ferric state to modulate its autokinase activity resulting in increased phosphorylation of DosR. The ability of sulfide to bind DosS ferric heme iron is clearly distinct from that of NO and CO, which bind DosS only when its heme iron is reduced (Fig. 7). Thus, DosS exhibits remarkable plasticity in sensing multiple gasotransmitters regardless of the oxidation state of the heme iron to ensure induction of the Dos dormancy regulon during infection. These findings establish a new paradigm for how bacteria sense multiple signaling molecules with distinct physicochemical properties through a single heme sensor kinase.

**Fig. 7.**
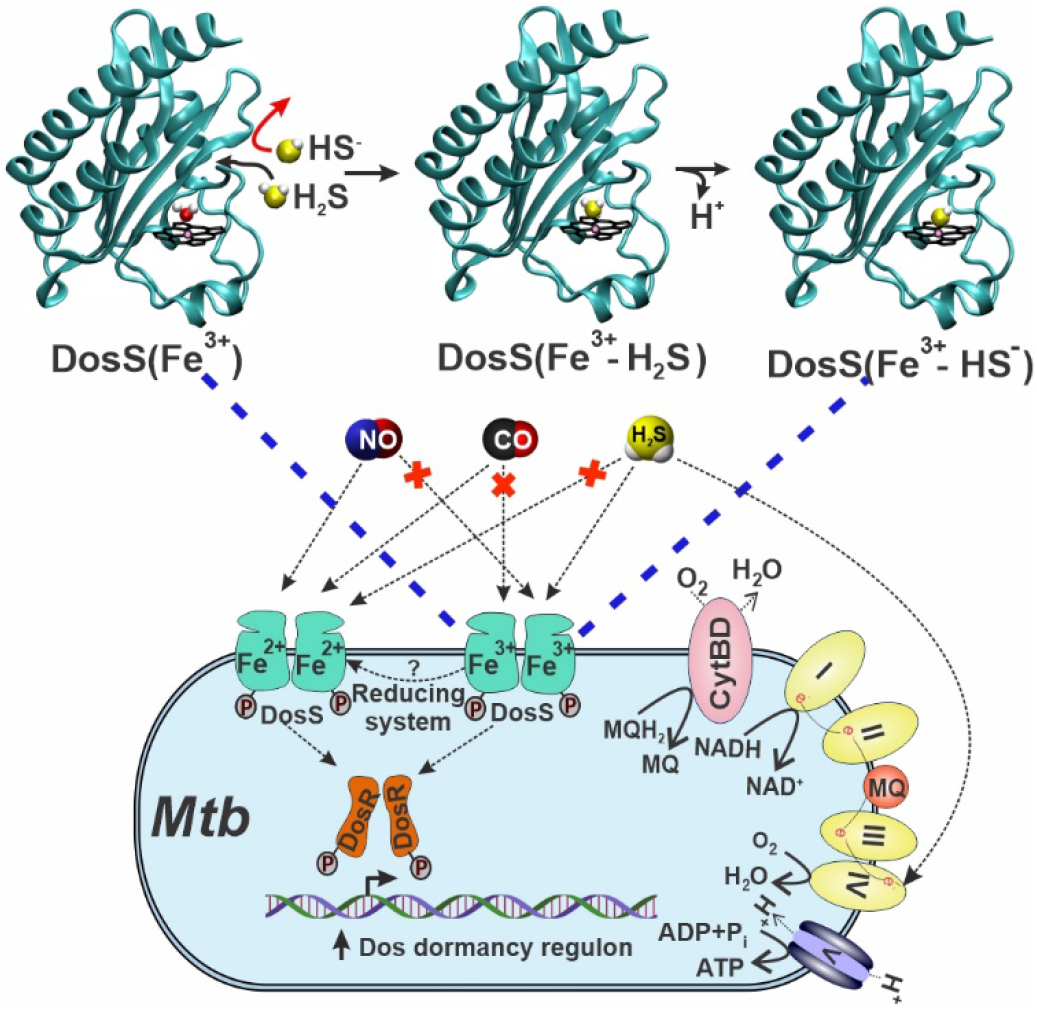
Schematic of a proposed mechanism of H_2_S sensing by *Mtb*. H_2_S, and not HS^-^ (red arrow), can enter the DosS heme pocket. Upon entering the heme pocket, H_2_S binds to DosS only in the ferric (Fe^3+^) state, demonstrating that sulfide sensing is different than for NO and CO, which require ferrous (Fe^2+^) DosS. Similar to NO and CO, binding of H_2_S to DosS increases autokinase activity which ultimately leads to increased expression of Dos dormancy regulon genes.

The importance of H_2_S as a signaling molecule in the regulation of numerous important physiological functions in mammals, including immunity, is well established (42, 43). Hence, it is reasonable that bacterial pathogens have evolved the capacity to detect H_2_S to reprogram their transcriptomes to subvert the immune response. However, while numerous signaling studies have been performed in bacteria including, but not limited to, *E. coli*, a gut organism continuously exposed to H_2_S, little is known regarding whether H_2_S has a direct signal transduction function in bacteria. More specifically, evidence supporting the ability of bacteria to sense and respond to host-derived H_2_S via heme sensor kinases in two-component systems is lacking (44, 45).

Of note, H_2_S is produced at the site of *Mtb* infection, as demonstrated by the presence of H_2_S-producing enzymes CSE and 3-MPST around cavities and necrotic lesions in the lungs of TB patients (6). Since H_2_S is highly diffusible, intracellular and extracellular *Mtb* will be exposed to host-generated H_2_S. *Mtb* is also exposed to NO (3), CO (2) and hypoxia (46) in the lungs of TB patients depending on disease stage. NO, CO and H_2_S are produced at different levels and times throughout the course of infection, which likely vary depending on the particular microanatomical location and pathology induced by *Mtb*. This, coupled with differing on- and off-rates of these molecules for DosS and DosT heme iron, strongly suggest that the *Mtb* Dos dormancy regulon is induced throughout the course of infection. Studies in the macaque model of inhalation TB have shown that *Mtb* Δ*dosS, ΔdosR* and Δ*dosT* mutants are attenuated (47). However, a *Mtb* Δ*dosS* mutant, but not Δ*dosR* or Δ*dosT* mutants, was shown to be attenuated in C3HeB/FeJ mice and macrophages (48). Notably, these authors demonstrated that DosS phosphorylates proteins other than DosR, suggesting that DosS may modulate expression of genes not in the Dos dormancy regulon (49). These data suggest that the mechanisms by which DosS senses and responds to environmental signals is more complex than previously thought.

Several lines of evidence provide insight into the mechanism whereby *Mtb* senses H_2_S. Firstly, we found that DosS, but not DosT, is a H_2_S sensor as the latter is in the ferrous oxy-bound state (Fe^2+^-O_2_) under normoxic conditions. Secondly, sulfide binds DosS under normoxic conditions where its heme iron is in the ferric form, as demonstrated by UV-Vis and EPR spectroscopy. Our spectroscopic data also show that binding of sulfide does not alter the oxidation state of DosS ferric heme iron, and that direct H_2_S binding to the ferric iron changes its spin state from high to low. Our findings are consistent with several reports showing that sulfide reacts with heme iron in the ferric (Fe^3+^) state (15-19). This may allow induction of the Dos regulon at earlier stages of infection prior to the onset of hypoxia when ferrous DosS can respond to NO and CO only. Thirdly, our MD modeling using the DosS GAF-A domain X-ray structure (PDB code 2W3E) (22) provides key insight into the molecular interactions of sulfide with DosS. Consistent with our sMD modeling of other hemoproteins (27, 28) H_2_S has more favorable access to the DosS heme than does HS^-^, which can explain the increase in H_2_S binding to DosS with decreasing pH. This selectivity is due to the strong hydration of HS^-^ compared to H_2_S which significantly hinders HS^-^ entry into the hydrophobic pocket, particularly *via* a “gate” comprised of Phe^98^ and Leu^114^ side chains. Further, QM[DFT]/MM calculations demonstrate a stronger Fe-S bond for HS^-^ than for H_2_S, suggesting deprotonation is induced upon binding of H_2_S. Indeed, our hybrid QM/MM simulations indicate proton transfer from H_2_S via a water molecule to a heme propionate resulting in bound HS^-^ as the final state.

To the best of our knowledge, this is the first report of a heme sensor kinase that binds H_2_S to increase its catalytic activity under physiologically relevant conditions. An important remaining question is how the binding of H_2_S to DosS increases autokinase activity. Given that a complete crystal structure for DosS has not been reported, a detailed study of the structural changes in the kinase domain upon H_2_S binding remains challenging. Nonetheless, structural studies probing the mechanism by which DosS activity is modulated by the redox state of, and ligand binding to, the heme iron have shown that an intact hydrogen bonding network comprised of Tyr^171^-Glu^87^-His^89^ is important for inhibition of DosS kinase activity, as seen in the Fe^3+^ state (22). Disruption of this network upon reduction of the heme iron and subsequent binding of NO or CO result in DosS conformations that favor kinase activity (22, 26, 50, 51). Our MD- and QM/MM-based modeling of the DosS GAF-A domain predicts that HS^-^ ligation to DosS disrupts the distal Tyr^171^-Glu^87^-His^89^ hydrogen bonding network. This is consistent with previously published proposals regarding signal transmission between the heme and autokinase domains (26) and with our finding of H_2_S-stimulated autokinase and DosR phosphorylation.

Our finding that H_2_S binds DosS in the Fe^3+^ state only, and not DosT which is in the Fe^2+^-O_2_ state, demonstrates that the mechanism of H_2_S sensing is distinct from NO and CO, since these molecules bind DosS in the Fe^2+^ state only. We (10) and others (22, 52) have shown that DosS exists in the Fe^3+^ state and requires a reductant to generate ferrous DosS that binds NO or CO. Ferredoxin (FdxA), reduced flavin nucleotides (22), and chorismate synthase (CS) have been posited as reductants of ferric DosS (22, 53, 54). FdxA and CS may be candidate reductants as *fdxA* is part of Dos regulon which is upregulated under hypoxic conditions and CS accelerates NADH-dependent FMN reduction. These potential reducing systems for DosS-Fe^3+^ may represent an additional aspect of Dos regulon induction, and suggests that in the absence of a reducing system, the Fe^3+^ form of DosS can still induce the Dos regulon via binding of H_2_S. Hence, in the presence of NO, CO or H_2_S, the Dos regulon will be induced via DosS regardless of whether the heme iron is in the Fe^2+^ or Fe^3+^ (unligated or ligated) state. Thus, these findings suggest that DosS signaling arising due to its redox sensor function and ligand binding are equally important. This may have important consequences *in vivo* and again suggests that the Dos dormancy regulon is induced through most of the course of infection. It has been shown that NO and hypoxia (11, 55), and likely CO, inhibit respiration, and all three factors induce the Dos regulon (5, 11, 55). However, we have shown that H_2_S increases *Mtb* respiration at low concentrations (7) and induces the Dos regulon under normoxic conditions (Fig. 6 and (7)). These findings suggest that inhibition of respiration (or hypoxia) is not the sole factor that induces the Dos dormancy regulon, and are consistent with the action of the TB drug bedaquiline, which also stimulates respiration (56) and induces the Dos dormancy regulon (57).

We (Fig. 1B, (10)) and others (22, 58) have shown that purified DosS exists in the Fe^3+^ state; however, others have reported DosS to be in the ferrous-oxy bound (Fe^2+^-O_2_) form (20, 54). The reasons for this discrepancy remain unclear but are likely due to differences in protein purification methodology and experimental approaches. Nonetheless, our EPR and UV-Vis spectroscopy data spectroscopy provide compelling evidence that DosS heme is in the ferric state (Fig 1B and 1D). Also, our data showing that DosS can only bind H_2_S in the met (Fe^3+^) state is consistent with numerous studies demonstrating that ferric, but not ferrous heme iron in proteins binds H_2_S (15, 16, 18, 45).

In light of previous studies showing that *Mtb* infection of macrophages induces upregulation of host H_2_S-producing enzymes CBS (7) and CSE (6), which leads to excessive levels of H_2_S to exacerbate disease, it was important to demonstrate that *Mtb* is capable of sensing physiological levels of H_2_S. Indeed, we demonstrated that WT *Mtb*, but not *Mtb ΔdosS*, senses endogenous levels of H_2_S during infection to induce key genes in the Dos regulon *fdxA, hspX, rv2030c*, and *rv2626*. This was further confirmed by enhancing H_2_S production in macrophages via substrate supplementation (6), which lead to increased expression of these Dos dormancy genes. This provides strong evidence that DosS senses H_2_S during infection.

To the best of our knowledge, this is the first report of a heme sensor kinase that binds H_2_S to increase its catalytic activity under physiologically relevant conditions. Further, our results show that H_2_S can function as a signaling molecule in *Mtb*. An important remaining question is how the binding of H_2_S to DosS increases autokinase activity. Given that a complete crystal structure for DosS has not been reported, a detailed study of the structural changes in the kinase domain upon H_2_S binding remains challenging. However, our MD-based modeling predicts that H_2_S binding to DosS disrupts the hydrogen bonding network in the distal domain, which is thought to be a mechanism of DosS activation (26, 50, 52).

In summary, we have shown that H_2_S binds to the redox sensor DosS to increase its autokinase and phosphate transfer activity, which induces the Dos dormancy regulon (Fig. 7). We have also shown that physiological levels of H_2_S are sufficient to induce the Dos regulon via DosS. The ability of *Mtb* to induce the Dos regulon in response to four physiologically relevant gasotransmitters points to a sophisticated signal transduction system to ensure *Mtb* persistence.

## Materials and Methods

### Mycobacterial strains and culturing conditions

*Mtb* H37Rv (BEI Resources (NR-123) and the *Mtb* H37Rv deletion mutant *ΔdosS* (provided by Dr. David Sherman, University of Washington) were grown at 37 °C with shaking in Middlebrook 7H9 medium (Difco) supplemented with 10% (vol/vol) ADS (albumin, dextrose, sodium chloride), 0.2% glycerol and 0.02% tyloxapol. Sodium sulfide (Na_2_S.9H_2_O) (Sigma) was added to culture media as an H_2_S donor. Stock solutions of Na_2_S were prepared in argon-deoxygenated 50 mM sodium phosphate buffer (150 mM NaCl, 5% glycerol, pH 7.4) and contained 100 µM DTPA (Diethylenetriaminepentaacetic acid) as a metal chelator.

### Expression and Purification of Recombinant DosS and DosT

DosS and DosT were expressed in Rosetta (DE3) BL21 *E*.*coli* cells grown at 37 °C in LB medium as reported previously (10). Briefly, bacterial cells were grown at 37 °C to an OD_600_ of 0.5-0.6, at which point hemin (Sigma) was added to a final concentration of 20 µM hemin and protein production was induced by addition of IPTG to a final concentration of 0.4 mM. Cells were grown overnight at 18°C, collected by centrifugation, and lysed by sonication in sodium phosphate buffer (pH 7.4). Soluble proteins were extracted by using Profinity IMAC Ni-charged resin (Bio-Rad, Hercules, CA) as recommended by the manufacturer. Following elution, proteins were dialyzed against sodium phosphate buffer (pH 7.4) to remove imidazole.

### UV-Vis Absorption Spectroscopy

The absorbance spectra of recombinant DosS and DosT (3 µM) were determined at room temperature using quartz cuvettes with rubber stoppers in a DU800 spectrophotometer (Beckman Coulter, Fullerton, CA). The reduction of recombinant DosS was achieved via addition of sodium dithionite (DTH) to a final concentration of 100 µM while inside an anaerobic glove box (Plas-Labs, Inc. Lansing, MI). A Hamilton syringe with a thin-gauge needle was used to add Na_2_S to protein solutions contained in quartz cuvettes (Spectrocell) sealed with a screw cap containing a rubber septum.

### EPR Spectroscopy

Purified DosS protein (10 µM) in sodium phosphate buffer with and without Na_2_S was transferred to thin-walled quartz EPR sample tubes (Wilmad Glass, Buena, NJ) and snap-frozen in liquid nitrogen. Cryogenic (7K) EPR was measured on a Bruker EMX spectrometer operating at a frequency of 9.39 GHz, 15-G modulation amplitude, 33 db power, 81.92-ms time constant, and 41.94-s sweep time. Each sample was scanned 4-8 times and the average taken.

### Determination of K_D_^p^for H_2_S binding to DosS

The K_D_^app^ for H_2_S binding to DosS was determined by difference spectroscopy using the Soret region. Fractional saturation was determined assuming complete occupancy at 100 µM [H_2_S] + [HS^-^]; this was verified by comparing to 200 µM.

### *In Vitro* Phosphorylation Assay

*In vitro* autokinase assays were performed essentially as described (33). Briefly, recombinant DosS (6 µM) alone or in the presence of 100 μM Na_2_S was assayed for its ability to autophosphorylate in a reaction containing 50 µCi of [*γ*-^32^P]-labeled ATP (6000 Ci/mmol, PerkinElmer Health Sciences), 100 mM Tris-HCl, pH 7.4, 5 mM MgCl_2_, 50 mM KCl_2_ in a final volume of 20 µl. The reaction was carried out at room temperature, and 4 µl aliquots of reaction mixture were removed from the reaction at various time points between 0 to 60 mins. The reaction at each time point was stopped by adding 2X SDS-PAGE sample buffer. The samples were resolved on 4-20% gradient PAGE gels (Bio-Rad) without heating. Resolved proteins were transferred to a PVDF membrane which was exposed to a phosphor screen (Amersham) overnight. The phosphor screen was scanned on a FLA7000IP Typhoon Storage Phosphorimager. Transphosphorylation assays were performed as above, except that DosS and DosR proteins were present in a molar ratio of 1:6, respectively.

### *Mtb* RNA Isolation and qRT-PCR Analysis

*Mtb* was grown to an OD_600_ of 0.4-0.5, and then Na_2_S was added to a final concentration of 50 µM. After incubation at 37 °C for 30 min, a volume of 4M guanidine thiocyanate solution equal to the culture volume was added, and cells were collected by centrifugation. The cell pellet was washed 1X time with PBS and then suspended in 1 ml RNAPro solution (MP Biomedicals). The cells were then added to a 2 mL tube containing 0.1 mm Lysing Matrix B beads (MP Biomedicals) and were lysed using a Fastprep-24 bead beater(MP Biomedicals). The lysate was centrifuged to pellet cell debris. 500 μl of chloroform was added to the supernatant fraction which was then vortexed and centrifuged for phase separation. Total RNA was isolated from the aqueous layer using a Qiagen RNA isolation kit, following the manufacturer’s instructions. RNA was treated with DNaseI prior to qRT-PCR analysis. 500 ng of DNaseI-treated RNA was used for cDNA synthesis using the iScript cDNA synthesis kit (Bio-Rad, USA). Quantitative RT-PCR was performed using SsoAdvanced SYBR green supermix (Bio-Rad, USA) with the Bio-Rad CFX96 detection system according to the manufacturer’s instructions. Quantitative RT-PCR reactions were performed in duplicate using three independent *Mtb* cultures. Relative changes in gene expression were determined using the 2ΔCt method (59), where Ct values of target genes were normalized to Ct values of *Mtb sigA* mRNA. Relative changes were determined as the ratio of expression of genes between untreated and Na_2_S-exposed cultures. Primers used in this study are listed in Table 1.

### Macrophage infection and isolation of intracellular *Mtb*

RAW 264.7 macrophages (ATCC) were cultured in DMEM medium (Gibco) containing 10% heat-inactivated FBS and 10 mM HEPES to maintain pH 7.4. RAW 264.7 macrophages were grown in 75 cm^2^ flasks and infected at a MOI of 1:10 for 2 hours using *Mtb* in log-phase growth. After removing infection media, cells were washed 2X with fresh DMEM medium and incubated in complete media containing the desired concentration of L-cysteine (0-2.0 mM) for 24 hours. The L-cysteine-containing media was removed and cells were washed 1X time with fresh media. Next, intracellular *Mtb* bacilli were isolated essentially as described by Rohde et al. (60) with a few modifications. Briefly, infected macrophages were lysed in a solution of 4M guanidine thiocyanate, 0.5% Na N-lauryl sarcosine, 25 mM sodium citrate, and 0.1 M β-mercaptoethanol. Lysate were vortexed and passed through a 21 gauge needle ten times to shear genomic DNA and reduce viscosity. Intracellular mycobacteria were recovered by centrifugation at ∼2,700 x g for 30 min. The bacterial pellet was suspended in 100 µl PBS and lysozyme was added to a final concentration of 0.1 mg/ml an incubated for 30 min at RT. The bacilli were lysed in 1 ml Trizol heated to 65°C with 0.1 mm Lysing Matrix B silicon beads using a Fastprep-24. Next, the lysate was centrifuged to pellet cell debris and the supernatant fraction was treated with 500 μl of chloroform, vortexed, and centrifuged. RNA was isolated from the aqueous layer using a Qiagen RNA isolation kit, following the manufacturer’s instructions. RNA was DNase treated prior to qRT-PCR analysis. A negative control group containing bacteria only was treated with complete DMEM was used to determine baseline gene expression levels.

### Detection of Intracellular H_2_S Using the WSP-5 Probe

The H_2_S-specfic WSP-5 fluorescent probe (CAS # 1593024-78-2) was prepared according to the manufacturer’s instructions. Cell staining was performed as described (36) with some modifications. Briefly, 5×10^4^ RAW 264.7 macrophages were plated in each well of 96 well plate and incubated overnight in DMEM medium containing the desired concentration of L-cysteine. The next day the medium was removed, and fresh medium containing 100 µM CTAB, the desired L-cysteine concentration, and 50 μM WSP-5 probe was added. After 30 min, the WSP-5-containing medium was removed, the cells were washed once with PBS and placed in fresh PBS for imaging. Fluorescence imaging was performed using Biotek Cytation 5 plate reader.

## Computational Methods

### Starting Structure

The starting structure of the DosS heme-containing GAF-A domain was built from the corresponding X-ray structure (PDB entry code 2w3e) (22). Protonation states of amino acids were those corresponding to their physiological state at neutral pH (i.e., Asp and Glu negatively charged, Lys and Arg positively charged). His149, corresponding to the heme proximal ligand, was simulated in the HID state (with protonated N δ). The remaining His residues were simulated favoring H-bond formation. Particular care was taken for His^89^ which is part of the distal H-bond network. Since the template crystal structure consists of a truncated GAF-A domain, a C-terminal carboxyl group is present that is not present in the full structure. To account for this, a N-CH_3_ “cap” was added to this carboxyl groups to avoid interaction with nearby positive residues. The system was solvated by constructing an octahedral box of 10-12 Å using AmberTools.

### Classical Molecular Dynamics (MD) Simulations

The MD simulations were performed using the Assisted Model Building with Energy Refinement (AMBER) package (61). Heme, as well as bound and free H_2_S and HS^-^ parameters, were taken from our previous work related to H_2_S/HS^-^ binding to heme proteins (28). All MD simulations were performed using periodic boundary conditions, SHAKE algorithm and the particle mesh Ewald (PME) summation method for treating the electrostatic interactions using default AMBER 16 parameters (61). Each system was first optimized, and then slowly equilibrated to reach proper temperature and pressure values using the Langevin thermostat and Berendsen barostat (62). All four truncated endpoints of the protein (exposed to solvent) were simulated with mild restraints applied to them to maintain α-helix structure.

### Ligand Binding Free Energy Profiles

To determine H_2_S/HS^-^ binding free energy profiles, we used our previously developed and extensively used Steered MD (sMD) approach combined with Jarzynski’s equation (63, 64). Briefly, in each simulation the ligand is pulled towards the heme iron inside the protein active site using a harmonic guiding potential at constant speed, and the corresponding work performed on the system is recorded. Several simulations are performed starting from different initial conformations with the ligand outside the protein, and finally the corresponding free energy profile is obtained combining the corresponding work profiles using Jarzynski’s equality. We employed 98 different trajectories for H_2_S and 89 for HS^-^. In all cases, the guiding coordinate was the Fe-S distance, using a force constant of 200 kcal/mol.Å^2^ and a speed of 0.0025 Å/ps.

### Hybrid Quantum Mechanics/Molecular Mechanics Simulations

QM/MM simulations were performed with our own extensively tested and developed code, called LIO which combines a Gaussian-based density functional theory (DFT) approach implemented in CUDA with the AMBER force field and is implemented as a QM/MM option in AMBER (65). System parameters were the same as in our previous QM/MM work on heme proteins (26, 28). Briefly, the heme, its proximal and distal ligands, a nearby water molecule and a heme propionate define the QM subsystem, and the remaining protein and solvent atoms the classical system. Covalent bonds between QM and MM subsystems were treated using the Link atom method (66). The QM system is simulated using a double Z plus polarization basis set (67) and the PBE exchange-correlation functional (68). Heme iron in the Fe^3+^ state bound to H_2_S/HS^-^ was simulated in the low-spin state. We performed a 2.0 ps MD simulation, starting from a snapshot extracted from the previously optimized MD simulation.

## Acknowledgements

This work was supported by NIH grants R01AI111940, R01AI138280, R01AI134810 (AJCS) and R01HL098032 (DBKS), a Bill and Melinda Gates Foundation Award (OPP1130017) (AJCS) and pilot funds from the UAB CFAR, CFRB, and Infectious Diseases and Global Health and Vaccines Initiative (AJCS). The research was also funded in part by the South African Medical Research Council to AJCS. QM/MM simulation studies were supported by grants from the Universidad de Buenos Aires (UBACYT 20020120300025BA), Agencia Nacional de Promoción Científica y Tecnológica (PICT 2016-0568, PICT 2014-1022, and PICT 2015-2761) and CONICET Grant 11220150100303CO. M.P. holds a CONICET Ph.D. fellowship. The authors wish to thank the staff and management of Southeastern Biosafety Laboratory Alabama Birmingham (SEBLAB), a NIAID-supported (UC6 AI058599) Regional Biocontainment Laboratory. The authors wish to thank Hayden T. Pacl for assistance in image analyses.

## Notes

**Competing Interest Statement:** The authors declare no conflicts of interest.

### Competing Interest Statement

The authors have declared no competing interest.

## References

1. Anonymous, Global tuberculosis report 2020. Geneva: World Health Organization; 2020. Licence: CC BY-NC-SA 3.0 IGO. (2020).

2. K. C. Chinta et al., Microanatomic Distribution of Myeloid Heme Oxygenase-1 Protects against Free Radical-Mediated Immunopathology in Human Tuberculosis. Cell Rep 25, 1938–1952 e1935 (2018).

3. S. Nicholson et al., Inducible nitric oxide synthase in pulmonary alveolar macrophages from patients with tuberculosis. J Exp Med 183, 2293–2302 (1996).

4. A. Kumar et al., Heme oxygenase-1-derived carbon monoxide induces the Mycobacterium tuberculosis dormancy regulon. J Biol Chem 283, 18032–18039 (2008).

5. M. U. Shiloh, P. Manzanillo, J. S. Cox, Mycobacterium tuberculosis senses host-derived carbon monoxide during macrophage infection. Cell Host Microbe 3, 323–330 (2008).

6. M. A. Rahman et al., Hydrogen sulfide dysregulates the immune response by suppressing central carbon metabolism to promote tuberculosis. Proc Natl Acad Sci U S A 117, 6663–6674 (2020).

7. V. Saini et al., Hydrogen sulfide stimulates Mycobacterium tuberculosis respiration, growth and pathogenesis. Nat Commun 11, 557 (2020).

8. S. T. Cole et al., Deciphering the biology of Mycobacterium tuberculosis from the complete genome sequence. Nature 393, 537–544 (1998).

9. A. K. Kinger, J. S. Tyagi, Identification and cloning of genes differentially expressed in the virulent strain of Mycobacterium tuberculosis. Gene 131, 113–117 (1993).

10. A. Kumar, J. C. Toledo, R. P. Patel, J. R. Lancaster, Jr., A. J. Steyn, Mycobacterium tuberculosis DosS is a redox sensor and DosT is a hypoxia sensor. Proc Natl Acad Sci U S A 104, 11568–11573 (2007).

11. M. I. Voskuil et al., Inhibition of respiration by nitric oxide induces a Mycobacterium tuberculosis dormancy program. J Exp Med 198, 705–713 (2003).

12. L. Fu et al., Direct Proteomic Mapping of Cysteine Persulfidation. Antioxid Redox Signal 33, 1061–1076 (2020).

13. M. R. Filipovic, J. Zivanovic, B. Alvarez, R. Banerjee, Chemical Biology of H2S Signaling through Persulfidation. Chem Rev 118, 1253–1337 (2018).

14. R. Kumar, R. Banerjee, Regulation of the redox metabolome and thiol proteome by hydrogen sulfide. Crit Rev Biochem Mol Biol 10.1080/10409238.2021.1893641, 1–15 (2021).

15. F. R. S. D. Keilin, On the combination of Methaemoglobin with H_2_S. Proceedings of the Royal Society of London 113, 11 (1933).

16. B. Jensen, A. Fago, Reactions of ferric hemoglobin and myoglobin with hydrogen sulfide under physiological conditions. J Inorg Biochem 182, 133–140 (2018).

17. D. W. Kraus, J. B. Wittenberg, J. F. Lu, J. Peisach, Hemoglobins of the Lucina pectinata/bacteria symbiosis. II. An electron paramagnetic resonance and optical spectral study of the ferric proteins. J Biol Chem 265, 16054–16059 (1990).

18. A. C. Mot et al., Fe(III) -Sulfide interaction in globins: Characterization and quest for a putative Fe(IV)-sulfide species. J Inorg Biochem 179, 32–39 (2018).

19. R. Pietri et al., Factors controlling the reactivity of hydrogen sulfide with hemeproteins. Biochemistry 48, 4881–4894 (2009).

20. E. H. Sousa, J. R. Tuckerman, G. Gonzalez, M. A. Gilles-Gonzalez, DosT and DevS are oxygen-switched kinases in Mycobacterium tuberculosis. Protein Sci 16, 1708–1719 (2007).

21. R. J. P. W. D. W. Smith, The spectra of ferric haems and haemoproteins, Structure and Bonding (1970), vol. 7.

22. H. Y. Cho, H. J. Cho, Y. M. Kim, J. I. Oh, B. S. Kang, Structural insight into the heme-based redox sensing by DosS from Mycobacterium tuberculosis. J Biol Chem 284, 13057–13067 (2009).

23. K. Y. C. a. J. C. Morris, Kinetics of Oxidation of Aqueous Sulfide by O2. Environmental Science & Technology 6, 9 (1972).

24. J. Peisach, W. E. Blumberg, S. Ogawa, E. A. Rachmilewitz, R. Oltzik, The effects of protein conformation on the heme symmetry in high spin ferric heme proteins as studied by electron paramagnetic resonance. J Biol Chem 246, 3342–3355 (1971).

25. H. R. D. Takashi Y, John S. Leigh, Jr. George H. Reed, Michael R. Water-Man, and Toshio Asakura, Electromagnetic Properties of Hemoproteins. The Journal of Biological Chemistry 245, 2998–3003 (1970).

26. Y. Madrona, C. A. Waddling, P. R. Ortiz de Montellano, Crystal structures of the CO and NOBound DosS GAF-A domain and implications for DosS signaling in Mycobacterium tuberculosis. Arch Biochem Biophys 612, 1–8 (2016).

27. F. M. Boubeta et al., Mechanism of Sulfide Binding by Ferric Hemeproteins. Inorg Chem 57, 7591–7600 (2018).

28. F. M. Boubeta, S. E. Bari, D. A. Estrin, L. Boechi, Access and Binding of H2S to Hemeproteins: The Case of HbI of Lucina pectinata. J Phys Chem B 120, 9642–9653 (2016).

29. D. E. Bikiel et al., Modeling heme proteins using atomistic simulations. Phys Chem Chem Phys 8, 5611–5628 (2006).

30. L. Capece et al., Small ligand-globin interactions: reviewing lessons derived from computer simulation. Biochim Biophys Acta 1834, 1722–1738 (2013).

31. A. B. Anderson, C. R. Robertson, Absorption spectra indicate conformational alteration of myoglobin adsorbed on polydimethylsiloxane. Biophys J 68, 2091–2097 (1995).

32. H. Wajcman et al., Hemoglobin Redondo [beta 92(F8) His Asn]: an unstable hemoglobin variant associated with heme loss which occurs in two forms. Am J Hematol 38, 194–200 (1991).

33. D. M. Roberts, R. P. Liao, G. Wisedchaisri, W. G. Hol, D. R. Sherman, Two sensor kinases contribute to the hypoxic response of Mycobacterium tuberculosis. J Biol Chem 279, 23082–23087 (2004).

34. A. E. Braunstein, E. V. Goryachenkova, E. A. Tolosa, I. H. Willhardt, L. L. Yefremova, Specificity and some other properties of liver serine sulphhydrase: evidence for its identity with cystathionine -synthase. Biochim Biophys Acta 242, 247–260 (1971).

35. M. H. Stipanuk, P. W. Beck, Characterization of the enzymic capacity for cysteine desulphhydration in liver and kidney of the rat. Biochem J 206, 267–277 (1982).

36. B. Peng et al., Fluorescent probes based on nucleophilic substitution-cyclization for hydrogen sulfide detection and bioimaging. Chemistry 20, 1010–1016 (2014).

37. M. P. Brigham, W. H. Stein, S. Moore, The Concentrations of Cysteine and Cystine in Human Blood Plasma. J Clin Invest 39, 1633–1638 (1960).

38. G. Murphy et al., Prospective study of serum cysteine levels and oesophageal and gastric cancers in China. Gut 60, 618–623 (2011).

39. Y. V. Tcherkas, A. D. Denisenko, Simultaneous determination of several amino acids, including homocysteine, cysteine and glutamic acid, in human plasma by isocratic reversed-phase high-performance liquid chromatography with fluorimetric detection. J Chromatogr A 913, 309–313 (2001).

40. Y. Zhang et al., Dual emission channels for sensitive discrimination of Cys/Hcy and GSH in plasma and cells. Chem Commun (Camb) 51, 4245–4248 (2015).

41. H. D. Park et al., Rv3133c/dosR is a transcription factor that mediates the hypoxic response of Mycobacterium tuberculosis. Mol Microbiol 48, 833–843 (2003).

42. N. Dilek, A. Papapetropoulos, T. Toliver-Kinsky, C. Szabo, Hydrogen sulfide: An endogenous regulator of the immune system. Pharmacol Res 161, 105119 (2020).

43. R. Wang, Physiological implications of hydrogen sulfide: a whiff exploration that blossomed. Physiol Rev 92, 791–896 (2012).

44. V. Fojtikova et al., Effects of hydrogen sulfide on the heme coordination structure and catalytic activity of the globin-coupled oxygen sensor AfGcHK. Biometals 29, 715–729 (2016).

45. H. Takahashi et al., Hydrogen sulfide stimulates the catalytic activity of a heme-regulated phosphodiesterase from Escherichia coli (Ec DOS). J Inorg Biochem 109, 66–71 (2012).

46. M. Belton et al., Hypoxia and tissue destruction in pulmonary TB. Thorax 71, 1145–1153 (2016).

47. S. Mehra et al., The DosR Regulon Modulates Adaptive Immunity and Is Essential for Mycobacterium tuberculosis Persistence. Am J Respir Crit Care Med 191, 1185–1196 (2015).

48. U. S. Gautam et al., DosS Is required for the complete virulence of mycobacterium tuberculosis in mice with classical granulomatous lesions. Am J Respir Cell Mol Biol 52, 708–716 (2015).

49. U. S. Gautam et al., Mycobacterium tuberculosis sensor kinase DosS modulates the autophagosome in a DosR-independent manner. Commun Biol 2, 349 (2019).

50. D. Basudhar et al., Distal Hydrogen-bonding Interactions in Ligand Sensing and Signaling by Mycobacterium tuberculosis DosS. J Biol Chem 291, 16100–16111 (2016).

51. E. T. Yukl, A. Ioanoviciu, M. M. Nakano, P. R. de Montellano, P. Moenne-Loccoz, A distal tyrosine residue is required for ligand discrimination in DevS from Mycobacterium tuberculosis. Biochemistry 47, 12532–12539 (2008).

52. H. Y. Cho, H. J. Cho, M. H. Kim, B. S. Kang, Blockage of the channel to heme by the E87 side chain in the GAF domain of Mycobacterium tuberculosis DosS confers the unique sensitivity of DosS to oxygen. FEBS Lett 585, 1873–1878 (2011).

53. F. Ely et al., The Mycobacterium tuberculosis Rv2540c DNA sequence encodes a bifunctional chorismate synthase. BMC Biochem 9, 13 (2008).

54. A. Ioanoviciu, Y. T. Meharenna, T. L. Poulos, P. R. Ortiz de Montellano, DevS oxy complex stability identifies this heme protein as a gas sensor in Mycobacterium tuberculosis dormancy. Biochemistry 48, 5839–5848 (2009).

55. M. I. Voskuil, K. C. Visconti, G. K. Schoolnik, Mycobacterium tuberculosis gene expression during adaptation to stationary phase and low-oxygen dormancy. Tuberculosis (Edinb) 84, 218–227 (2004).

56. J. S. Mackenzie et al., Bedaquiline reprograms central metabolism to reveal glycolytic vulnerability in Mycobacterium tuberculosis. Nat Commun 11, 6092 (2020).

57. A. Koul et al., Delayed bactericidal response of Mycobacterium tuberculosis to bedaquiline involves remodelling of bacterial metabolism. Nat Commun 5, 3369 (2014).

58. H. Zheng et al., Inhibitors of Mycobacterium tuberculosis DosRST signaling and persistence. Nat Chem Biol 13, 218–225 (2017).

59. T. D. Schmittgen, K. J. Livak, Analyzing real-time PCR data by the comparative C(T) method. Nat Protoc 3, 1101–1108 (2008).

60. K. H. Rohde, R. B. Abramovitch, D. G. Russell, Mycobacterium tuberculosis invasion of macrophages: linking bacterial gene expression to environmental cues. Cell Host Microbe 2, 352–364 (2007).

61. K. B. D.A. Case, I.Y. Ben-Shalom, S.R. Brozell, D.S. Cerutti, T.E. Cheatham, III, V.W.D. Cruzeiro, T.A. Darden, R.E. Duke, G. Giambasu, M.K. Gilson, H. Gohlke, A.W. Goetz, R. Harris, S. Izadi, S.A. Izmailov, K. Kasavajhala, A. Kovalenko, R. Krasny, T. Kurtzman, T.S. Lee, S. LeGrand, P. Li, C. Lin, J. Liu, T. Luchko, R. Luo, V. Man, K.M. Merz, Y. Miao, O. Mikhailovskii, G. Monard, H. Nguyen, A. Onufriev, F. Pan, S. Pantano, R. Qi, D.R. Roe, A. Roitberg, C. Sagui, S. Schott-Verdugo, J. Shen, C.L. Simmerling, N.R. Skrynnikov, J. Smith, J. Swails, R.C. Walker, J. Wang, L. Wilson, R.M. Wolf, X. Wu, Y. Xiong, Y. Xue, D.M. York and P.A. Kollman (2020) AMBER 2020.

62. H. J. C. P. Berendsen, J. P. M.; van Gunsteren, W. F.; DiNola, A.; Haak, J. R, Molecular dynamics with coupling to an external bath. The Journal of Chemical Physics 81 (1984).

63. M. Arrar et al., On the accurate estimation of free energies using the jarzynski equality. J Comput Chem 40, 688–696 (2019).

64. C. Jarzynski, Nonequilibrium Equality for Free Energy Differences. Physical Review Letters 78, 2690–2693 (1997).

65. J. P. Marcolongo et al., Chemical Reactivity and Spectroscopy Explored From QM/MM Molecular Dynamics Simulations Using the LIO Code. Front Chem 6, 70 (2018).

66. M. E. a. P. Tavan, A hybrid method for solutes in complex solvents: Density functional theory combined with empirical force fields. The Journal of Chemical Physics 110 (1999).

67. N. Godbout, D. R. Salahub, J. Andzelm, E. Wimmer, Optimization of Gaussian-type basis sets for local spin density functional calculations. Part I. Boron through neon, optimization technique and validation. Canadian Journal of Chemistry 70, 560–571 (1992).

68. J. P. Perdew, K. Burke, M. Ernzerhof, Generalized Gradient Approximation Made Simple. Phys Rev Lett 77, 3865–3868 (1996).

