## Supplemental data for "*Mycobacterium tuberculosis* DosS binds H_2_S through its Fe^3+^ heme iron to regulate the Dos dormancy regulon"

Professor, University of Alabama at Birmingham

Assistant Professor, University of Alabama at Birmingham

**This PDF file includes:**

Figures S1 to S3

Table S1

Legends for Movies S1-S3

**Other supplementary materials for this manuscript include the following:**

Movies S1-S3

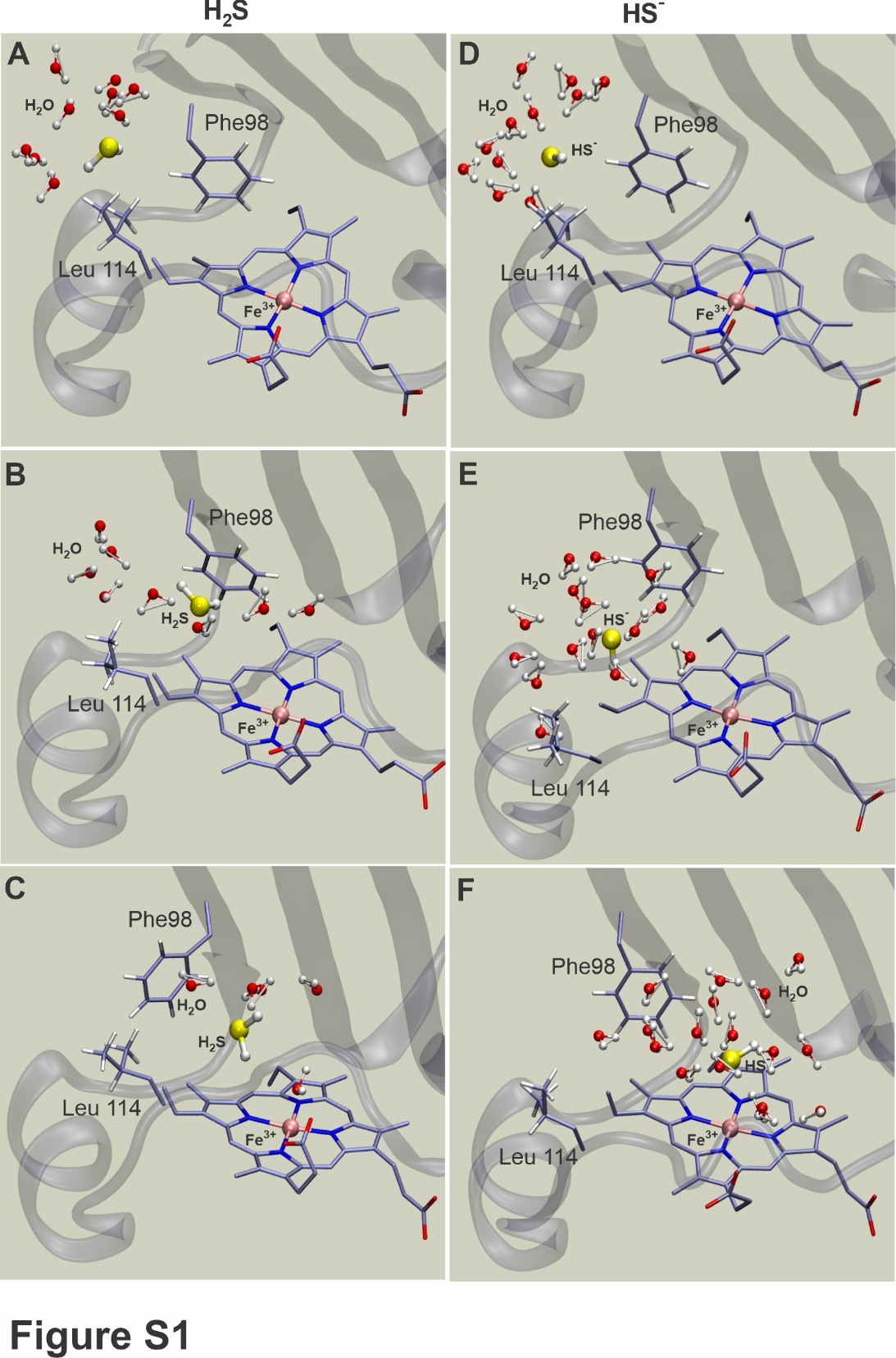

**Fig. S1.** **Steered Molecular Dynamics (sMD) modeling of** **sulfide entry into the DosS heme pocket.** Typical snapshots of H_2_S (A-C), and HS^-^ (D-F) along the entry path into the DosS heme pocket, representing sulfide ligands at a distance of 11, 8 and 5 Å (upper, medium, and lower panels, respectively) from the heme iron.

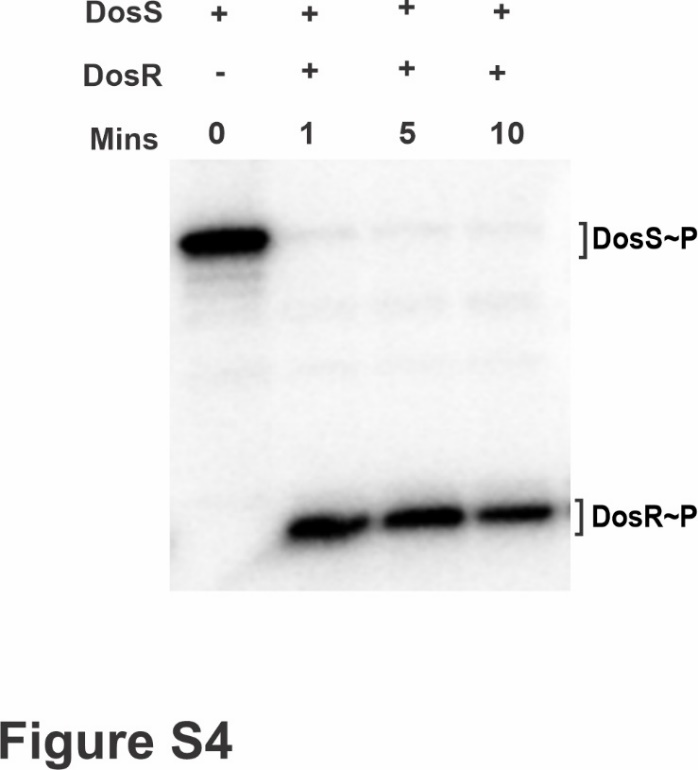

Fig. S2. Transphosphorylation assay. Autoradiogram of PAGE-resolved γ^32^P-labeled DosS in the Fe^3+^ form alone after 60 min of autophosphorylation (left lane), and following addition of recombinant DosR to the autokinase reaction. The reactions were stopped 1-10 min following addition of DosR.

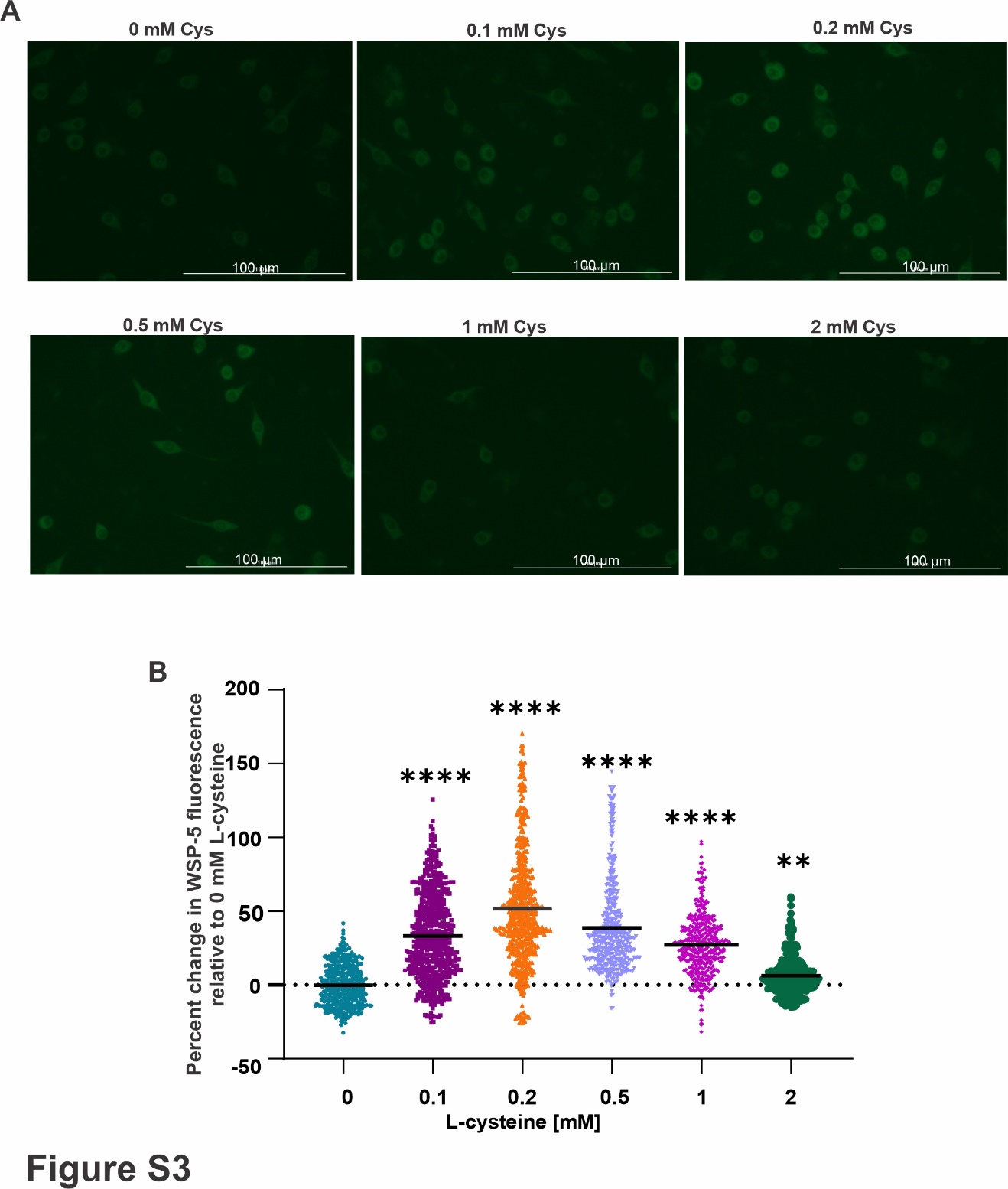

**Fig. S3**. **Detection of intracellular H_2_S.** (A) Representative images of intracellular WSP-5 fluorescence in RAW 264.7 macrophages grown in media containing 0-2.0 mM L-cysteine (Cys). (B) Percent change in intracellular H_2_S-dependent probe fluorescence levels in RAW 264.7 macrophages in presence of 0-2.0 mM L-cysteine as determined by WSP-5 fluorescence measurements in individual cells. The data were analyzed using Kruskal-Wallis test with Holm-Sidak multiple comparisons. The solid black line represents the mean value of the group. **P < 0.01 and ****P < 0.0001 vs 0 mM L-cysteine control.

**Table S1. DNA Primers used for qRT-PCR.**

| **Primer** | **Primer Sequence (5’ to 3’)** |
| --- | --- |
| SigA-For | TCGGTTCGCGCCTACCT |
| SigA-Rev | TGGCTAGCTCGACCTCTTCCT |
| hspX-For | CGCACCGAGCAGAAGGAC |
| hsp-Rev | CCGCCACCGACACAGTAA |
| fdxA-For | CCTATGTGATCGGTAGTGA |
| fdxA-Rev | GGGTTGATGTAGAGCATT |
| rv2030-For | GAATAGTGGTGTGGGCTCATAA |
| rv2030-Rev | CGTATCGCTCACGGACTATCT |
| rv2626-For | CGACCGCGACATTGTGAT |
| rv2626-Rev | CATCGACGTAGTAGATGCTGT |

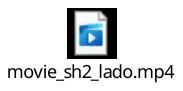

Movie S1. Steered Molecular Dynamics modeling of H_2_S entry into the DosS heme pocket.

The trajectory of H_2_S [shown in yellow (S) and white (H) spheres] approaching the DosS heme is shown. H_2_S first approaches the Y97-F98 pair, and then passes between F98 and the L114-P115 pair to enter the heme pocket, where it interacts with distal Y171. The H_2_S solvation shell waters are represented as small red and white triangles. The overall DosS secondary structure is shown as a cyan tube. Red = oxygen, Blue = nitrogen, Pink = heme center. Movie represents 2.8 ns of simulation with a ligand pulling speed of 0.0025 Â/ps.

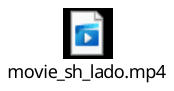

Movie S2. Steered Molecular Dynamics modeling of HS^-^ entry into the DosS heme pocket.

The trajectory of HS^-^ [shown in yellow (S) and white (H) spheres] approaching the DosS heme is shown. HS^-^ first approaches the Y97-F98 pair. The barrier for HS^-^ entry is formed by disrupting the F98-L114 interaction, which is clearly seen when the L114 sidechain changes separates from F98. HS^-^ can then pass between F98 and the L114-P115 pair to enter the heme pocket, where it interacts with distal Y171. The larger barrier for HS^-^ is due to the fact that while HS^-^ enters almost dry, the negatively charged ligand is always solvated and accompanied by several water molecules. The HS^-^ solvation shell waters are represented as small red and white triangles. The overall DosS secondary structure is shown as a cyan tube. Red = oxygen, Blue = nitrogen, Pink = heme center. Movie represents 2.8 ns of simulation with a ligand pulling speed of 0.0025 Â/ps.

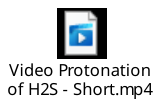

Movie S3. QM/MM modeling of deprotonation of H_2_S in DosS heme pocket.

Deprotonation of H_2_S [shown in yellow (S) and white (H) spheres] following binding to the DosS Fe^3+^ heme iron is shown. The proton transfer to a nearby heme propionate group via a water bridge is spontaneous and occurs within 0.35 ps. The QM subsystem is depicted in balls and sticks. The overall DosS secondary structure is shown as a gray ribbon. Red = oxygen, Blue = nitrogen, Pink = heme center. Movie represents 0.4 ns of simulation.

**SI References**

1.	D. W. Kraus, J. B. Wittenberg, J. F. Lu, J. Peisach, Hemoglobins of the Lucina pectinata/bacteria symbiosis. II. An electron paramagnetic resonance and optical spectral study of the ferric proteins. *J Biol Chem* **265**, 16054-16059 (1990).

2.	V. Vitvitsky, P. K. Yadav, A. Kurthen, R. Banerjee, Sulfide oxidation by a noncanonical pathway in red blood cells generates thiosulfate and polysulfides. *The Journal of biological chemistry* **290**, 8310-8320 (2015).

3.	A. C. Mot *et al.*, Fe(III) - Sulfide interaction in globins: Characterization and quest for a putative Fe(IV)-sulfide species. *J Inorg Biochem* **179**, 32-39 (2018).

4.	D. W. Kraus, J. B. Wittenberg, J. F. Lu, J. Peisach, Hemoglobins of the Lucina pectinata/bacteria symbiosis. II. An electron paramagnetic resonance and optical spectral study of the ferric proteins. *J.Biol.Chem* **265**, 16054-16059 (1990).

5.	T. Bostelaar *et al.*, Hydrogen Sulfide Oxidation by Myoglobin. *Journal of the American Chemical Society* **138**, 8476-8488 (2016).

6.	Z. Palinkas *et al.*, Interactions of hydrogen sulfide with myeloperoxidase. *Br J Pharmacol* **172**, 1516-1532 (2015).

7.	S. A. Bieza *et al.*, Reactivity of inorganic sulfide species toward a heme protein model. *Inorg Chem* **54**, 527-533 (2015).

8.	F. P. Nicoletti *et al.*, Sulfide binding properties of truncated hemoglobins. *Biochemistry* **49**, 2269-2278 (2010).

9.	H. Takahashi *et al.*, Hydrogen sulfide stimulates the catalytic activity of a heme-regulated phosphodiesterase from Escherichia coli (Ec DOS). *J Inorg Biochem* **109**, 66-71 (2012).

10.	V. Fojtikova *et al.*, Effects of hydrogen sulfide on the heme coordination structure and catalytic activity of the globin-coupled oxygen sensor AfGcHK. *Biometals* **29**, 715-729 (2016).
